# Spacer acquisition in type VI CRISPR-Cas systems associated with reverse transcriptase-Cas1 fusion proteins

**DOI:** 10.1101/2024.03.12.584598

**Authors:** María Dolores Molina-Sánchez, Francisco Martínez-Abarca, Vicenta Millán Casamayor, Mario Rodríguez Mestre, Nicolás Toro

**Affiliations:** Department of Soil and Plant Microbiology, Estación Experimental del Zaidín, Consejo Superior de Investigaciones Científicas, Structure, Dynamics and Function of Rhizobacterial Genomes, Grupo de Ecología Genética de la Rizosfera, C/Profesor Albareda 1, 18008, Granada, Spain

**Keywords:** Cas2, Cas13, CorA, RT-Cas1, spacer acquisition, type VI-A, type VI-B1

## Abstract

In prokaryotes, CRISPR-Cas systems store memories of past infections in the form of spacers integrated into CRISPR arrays. When associated with type III CRISPR-Cas systems, Reverse transcriptase-Cas1 fusion proteins (RT-Cas1) enable these defense systems to acquire spacers from RNA sources. However, despite the specific targeting of RNA by the Cas13-containing type VI CRISPR-Cas systems, there is no evidence of RNA-origin spacer acquisition. Using computational analyses, we recently reported the association of RT-Cas1 fusion proteins with type VI-A systems. In this study, we found that RT-Cas1 fusion proteins were also associated with complete type VI-B systems in bacteria from gut metagenomes, constituting a variant system that harbors a linked CorA-encoding locus in addition to the CRISPR array and adaptation RT-Cas1/Cas2 module. By combining *in vitro* and *in vivo* experiments, we demonstrated that type VI RT-CRISPR systems are functional for spacer acquisition and CRISPR array processing, and that the associated RT enables spacer acquisition from RNA molecules, thus demonstrating that the system is capable of functioning independently of other in-*trans* systems. These findings highlight the importance of RTs in RNA-targeting CRISPR-Cas systems, suggesting a potential defense mechanism against RNA-based invaders in specific environments.

## INTRODUCTION

The reverse transcriptases (RTs) of prokaryotes are highly diverse, and recent research has revealed an important role for these enzymes in defense mechanisms against phages and other mobile genetic elements. The RTs involved in defense are classified into various lineages, including retrons, Abi-like RTs, RTs of the so-called “unknown group” (UG), and those associated with CRISPR-Cas systems (Toro et al., 2019a, Mestre et al., 2020, González-Delgado et al., 2021, Mestre et al., 2022).

CRISPR-Cas (clustered regularly interspaced short palindromic repeats and CRISPR-associated proteins) systems are adaptive immune systems found in archaea and bacteria that can integrate fragments of invasive nucleic acids into CRISPR arrays, known as spacers, through a process called adaptation, which requires the Cas1/Cas2 proteins. The CRISPR array is transcribed and processed into short RNAs (crRNAs) that serve as guides for CRISPR-associated (Cas) proteins, enabling them to recognize and eliminate invasive DNA through complementarity. CRISPR-Cas systems can be classified in two classes, six types and 33 subtypes (Makarova et al. 2020). Type III CRISPR-Cas system are the only ones able to recognize and cleave both DNA and RNA targets in a transcription-coupled reaction (Samai et al., 2015; Marraffini 2022), whereas type VI are unique in their specific targeting of RNA molecules (Shmakov et al., 2015; East-Seletsky et al., 2016; Abudayyeh et al., 2016). The RT loci associated with CRISPR-Cas systems are either adjacent to or fused to the *cas1* gene (RT-Cas1), and most are linked to a subset of type III CRISPR-Cas systems (Silas et al. 2017; Toro et al., 2017; Toro et al. 2019a).

Phylogenetic analyses have suggested that most RTs associated with CRISPR-Cas adaptation modules probably evolved from the RTs encoded by group II introns (Silas et al. 2017, Toro et al. 2018, Toro et al., 2019a) after retrotransposition and subsequent domestication of the encoded RTs. Independent studies have shown that type III CRISPR-associated RT systems can acquire new spacers directly from RNA *in vivo* in an RT-dependent manner (Silas et al., 2016; Schmidt et al., 2018; González-Delgado et al., 2019). Interestingly, we have also reported the recruitment of RTCas1 fusion proteins in certain types of VI-A CRISPR-Cas systems (Toro et al. 2019b).

Type VI CRISPR-Cas systems often lack Cas1/Cas2, which has led to suggestions that they may acquire spacers in *trans* (Shmakov et al., 2015; Smargon et al., 2017; Barrangou and Gersbach, 2017; Zhu et al., 2018). In addition, a type VI-B system has been shown to acquire spacers using the Cas1/Cas2 machinery from a type II-C system (Hoikkala et al., 2021). However, the adaptation process of type VI CRISPR systems has yet to be elucidated.

In this study, we investigated spacer acquisition in type-VI CRISPR-RT-associated systems. We identified type VI-B systems in bacteria by analyzing gut metagenomes and genome assemblies associated with RT-Cas1 fusions. In addition to the CRISPR array and adaptation module, these systems harbor a linked CorA-encoding locus. We found that these RT-CRISPR systems were fully functional for spacer acquisition. Specifically, we demonstrated that in type VI-B systems, the associated RT enabled spacer acquisition from RNA molecules. These findings highlight the crucial importance of the integration of RTs into CRISPR-Cas systems, especially those that target RNA, providing them with a potent defense mechanism against RNA-based threats in specialized environments.

## RESULTS

### Spacer acquisition in CRISPR type VI-A systems with adaptation modules including RT-Cas1 fusion proteins

We previously reported two different CRISPR type VI-A systems featuring adaptation modules containing RT-Cas1 fusion proteins (Toro et al., 2019b). We investigated the spacer acquisition process within these VI-A systems, by focusing specifically on CRISPR-Cas loci representative of the two subtypes (**Table S1**). We designated *Rhodovulum kholense* loci as Rk, representing the type VI-A/RT1 system, and investigated two examples of the type VI-A/RT2 system, *Eubacterium rectale* AF25-15 (Er) and *Eubacteriaceae* bacterium CHKCI004 (CH), as shown in **Figure 1A**.

**Figure 1.**
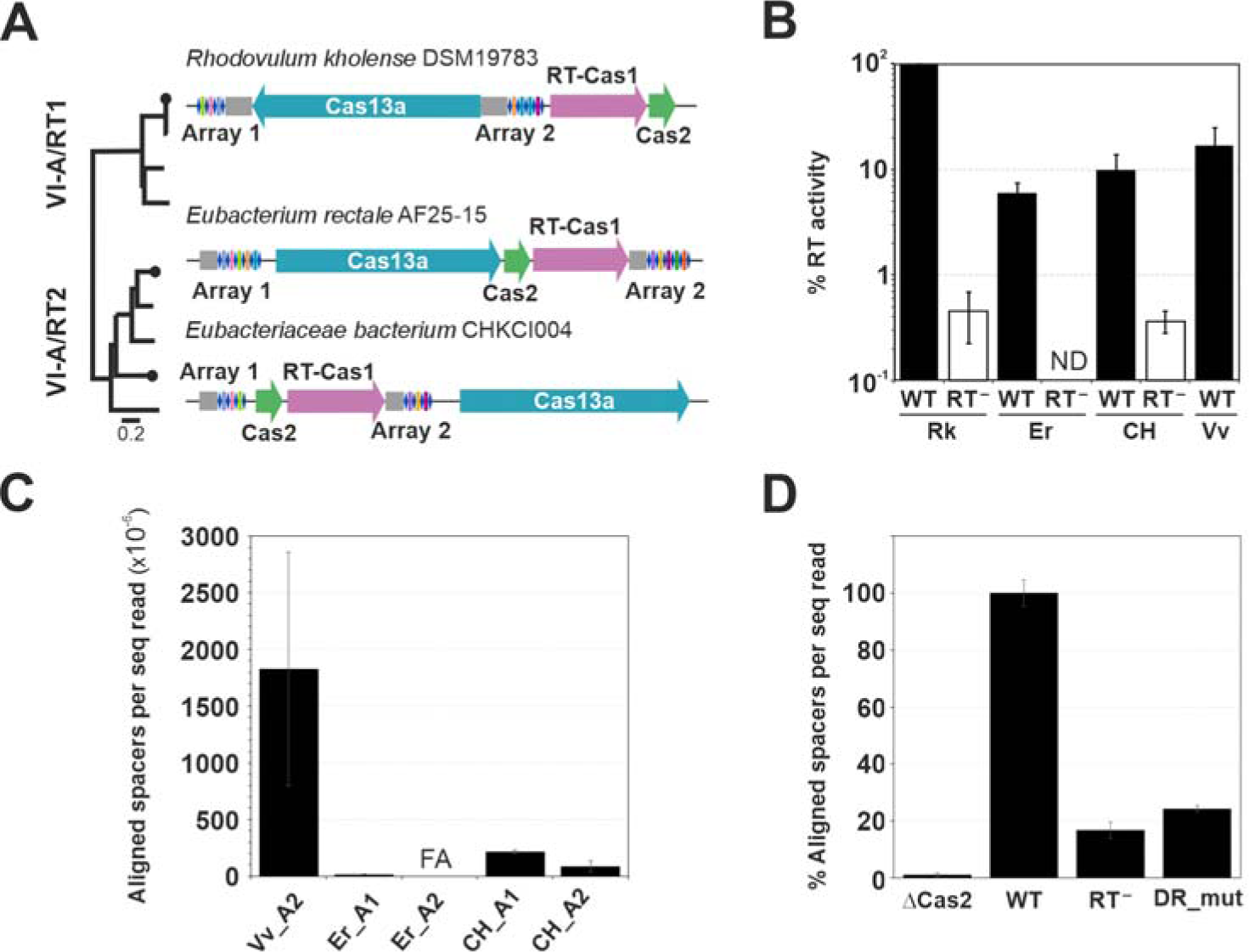
Acquisition of new spacers by CRISPR-Cas-type VI-A/RT systems. **(A)** The genomic arrangement of representative variants of the type VI-A/RT systems studied here is illustrated, with the phylogeny of RT-Cas1 proteins depicted on the left. Diagrams are drawn approximately to scale. Gray boxes indicate the predicted leader regions of arrays. *Rhodovulum kholense* DSM 19783, NZ_QAYC01000027.1; *Eubacterium rectale* AF25-15, NZ_QRUJ01000006.1; *Eubacteriaceae* bacterium CHKCI004, NZ_FCNR01000048.1 **(B)** Exogenous RT activity was measured by assessing the incorporation of [^32^P]-dTTP on a poly(rA) substrate primed with oligo(dT)_18_. Polymerization is presented as a percentage of the RT-Cas1 activity of Rk_WT, with the *y*-axis displayed on a logarithmic scale. The activity of the RT-Cas1 protein of a CRISPR-Cas Type III-D system from *Vibrio vulnificus* (Gonzalez-Delgado et al 2019) is included as a control. Black bars represent WT proteins, whereas white bars indicate RT-deficient mutant proteins. The data were derived from three technical replicates and at least one biological replicate and are presented as the mean ± standard error of the mean (SEM). Rk (*Rhodovulum kholense,* Er (*Eubacterium rectale*), CH (*Eubacteriaceae* bacterium), Vv (*Vibrio vulnificus*). **(C)** Spacer acquisition by type VI-A/RT systems in the heterologous host *E. coli* HMS174 (DE3). The frequency of newly acquired aligned spacers per million sequencing reads is shown for the wild-type adaptation operons from *E. rectale* (Er), and *Eubacteriaceae* CHKCI004 (CH), for both arrays (A1 and A2); the reported adaptation operon of *V. vulnificus* (Vv_A2) (Gonzalez-Delgado et al., 2019) was included as a control. Error bars represent the SEM and were calculated from at least three biological replicates (See methods). FA “Failed amplification” samples, for which the DNA yield from expanded arrays was too low for NGS. **(D)** Percentage of aligned spacers relative to the wild-type system of the *Eubacteriaceae* bacterium CHKCI004 on the A1 array (WT), normalized per sequencing read for the different mutants: ΔCas2 has no Cas2 protein gene in the adaptation module; RT-, the RT-Cas1 protein has a mutation in the catalytic core for RT activity; FGDD → FGAA; and DR_mut, the first two nucleotides of the array 1 DR are replaced with +1A +2C.

As we hypothesized that the type VI-A system-associated RT-Cas1 fusion protein might play a role in RNA spacer acquisition, we began by evaluating the ability of these RT-Cas1 fusion proteins to synthesize DNA from an RNA template. In our *in vitro* experiments, all maltose-binding protein (MBP)-RT-Cas1 fusion proteins tested displayed exogenous RT activity with poly (A)/oligo (dT) substrates. This RT activity was markedly higher than that of the corresponding RT-deficient mutants (**Figure 1B**). Furthermore, we found that mutations of critical residues of the Cas1 domain did not significantly disrupt DNA polymerization capacity (data not shown).

We then investigated whether either of the RT-Cas1-associated type VI-A systems studied could acquire new spacers from *E. coli,* used as a heterologous host. *E. coli* HMS174 (DE3) was transformed with a single plasmid expressing the adaptation module and the corresponding CRISPR array comprising the leader region, first DR and first natural spacer from a constitutive (pUC derivative) or inducible (pBAD backbone) promoter. Using the CAPTURE protocol with divergent primers (McKenzie et al., 2019; **Figure S1**), we subjected the PCR products to next-generation sequencing (NGS). For the Rk system, we detected no acquisition events with either the Rk_A1-derived or the Rk_A2 derived assay. However, in the Er system, new spacers were incorporated into the Er_A1 array, albeit through a highly inefficient acquisition process (**Figure 1C**). Spacer acquisition efficiency was greater for the CH system than for the Er adaptation module, but remained considerably lower than that of the *V. vulnificus* III-D/RT-Cas1 system (**Figure 1C**), as previously reported (González-Delgado et al., 2019). Importantly, we found that spacer acquisition efficiency was much lower in a mutant in which the direct repeat (DR) sequence was altered (with changes to the first and second residues of the DR, replaced by +1A and +2C, respectively). Similarly, both the RT-deficient mutant and the mutant lacking the Cas2 protein had much lower spacer acquisition efficiencies. These findings strongly suggest that new spacers are not acquired in *trans* using the acquisition machinery of other co-existing adaptive modules (**Figure 1D**).

In the CH type VI-A system, the most common spacer length was 34 bp for the chromosome (35 bp for the plasmid) (**Figure 2A**). An analysis of the GC content of the acquired spacers showed that it closely matched that of both the plasmid and the bacterial genome (**Figure 2B**). Moreover, examination of the protospacer regions flanking the aligned spacers revealed no sequence preference, suggesting that there was no protospacer adjacent motif (PAM) (**Figure 2C**). However, there was a tendency for the sequences surrounding both ends of the spacers to be more AT-rich.

**Figure 2.**
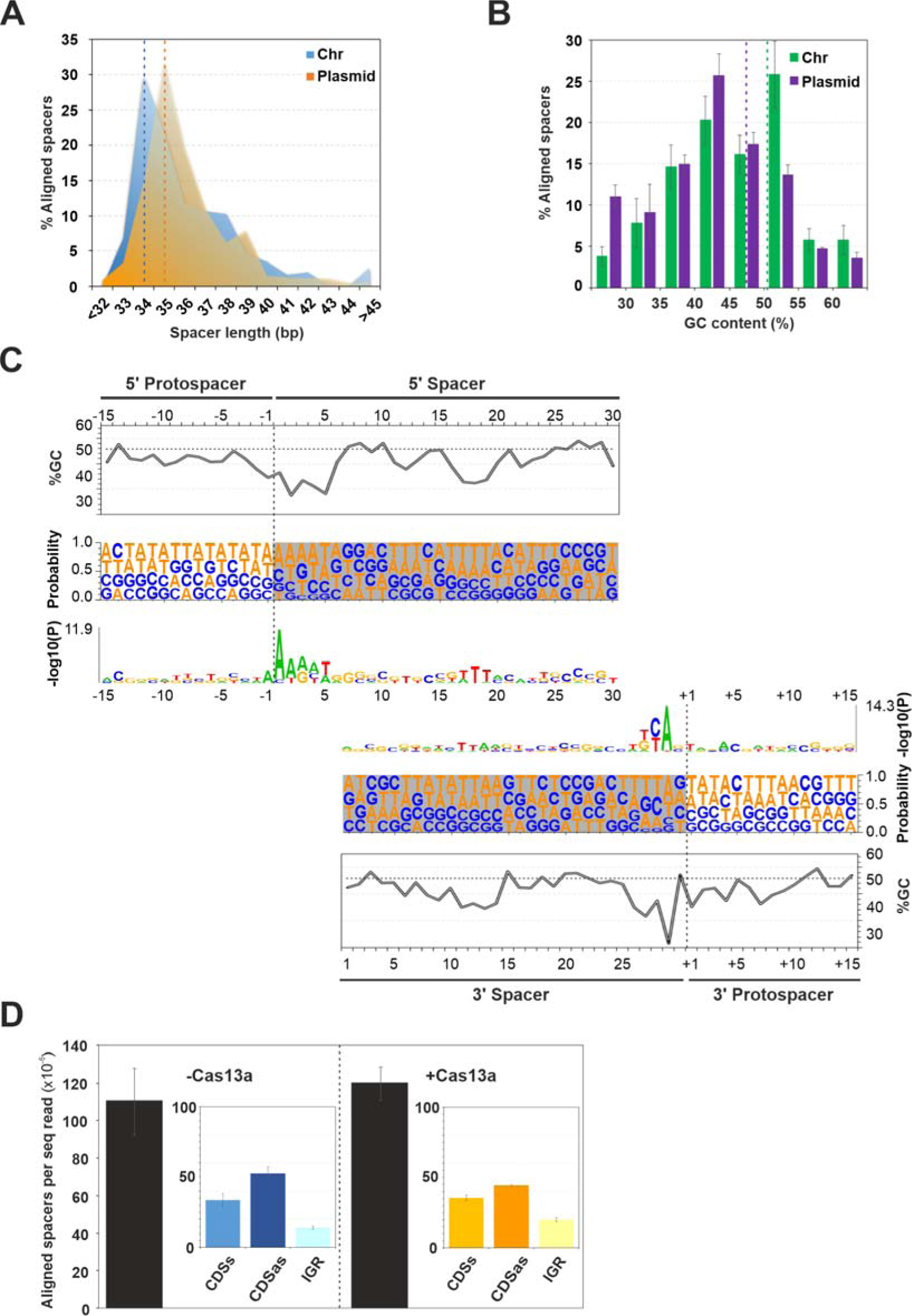
Characterization of spacers acquired by the *Eubacteriaceae* bacterium CHKCI004 adaptation system. **(A)** Distribution of newly detected spacers by length. Blue and orange indicate matches to the reads identified in the *E. coli* chromosome or the plasmid, respectively. **(B)** Distribution of GC content in the genome (green) and plasmid-aligned spacers (purple). The dotted lines represent the mean GC content of the plasmid (purple) and the genome (green). **(C)** Protospacer composition analysis for GC content, frequency and statistical probability. The spacer sequence is shaded and the spacer boundaries are marked as dotted lines. Owing to the different lengths of the detected spacers, the protospacer sequence was split into a 5’ protospacer (upper panels) and a 3’ protospacer (lower panels). Dotted lines in GC content plot correspond to the mean % GC in the *E. coli* genome. On the statistical probability graph [-log_10_ (P)], the size of the nucleotides represented indicates their relative probability (average across all positions) in the protospacer sequences analyzed according to k-mer length=1, 2, 3, 4 using a binomial test. All data (A-C) represent spacers merged across nine biological samples with the adaptation module in pUC. **(D)** Effect of Cas13a on spacer acquisition. Black bars represent aligned spacers detected per million sequencing reads in the absence (-) or presence (+) of Cas13a in pBAD-derived vectors. Inner boxes correspond to the proportion of newly acquired aligned spacers in the sense (CDSs) or antisense (CDSas) strand of coding genes or in intergenic regions (IGR) of the *E. coli* genome.

The activity of the conserved Cas1-Cas2 integrase complex is sufficient for type I CRISPR acquisition, but for type II-C CRISPR systems, the acquisition of new spacers also requires *cas9* and *csn2* (Yosef et al., 2012; Heler et al., 2015 and Wei et al., 2015). For the CH type VI-A/RT-Cas1 system, experiments in which the effector module (Cas13a) was added revealed no significant differences in the spacer acquisition efficiency or in the genomic regions used as a source of the acquired spacers in the presence or absence of this module, suggesting that the Cas13a effector may not be involved in the acquisition process (**Figure 2D**).

### Association of RT-Cas1 fusion proteins with CRISPR-Cas type VI-B systems

Given the low spacer acquisition efficiency observed for the type VI-A CRISPR-associated RT systems tested, we decided not to pursue our analysis of spacer acquisition from RNA sources. Instead, we conducted a comprehensive database analysis to identify additional type VI systems associated with RTs. We used the computational pipeline previously described by Toro et al. (2019a). Remarkably, within the non-redundant (nr) protein sequence database of the NCBI (https://www.ncbi.nlm.nih.gov/) and the MGnify (https://www.ebi.ac.uk/metagenomics) database, we were able to identify eight Cas13b effector proteins linked to adaptation modules featuring RT-Cas1 fusion proteins and Cas2 proteins. These proteins were identified exclusively from human, murine, and cockroach gut metagenomes and genome assemblies (**Table S1**), suggesting a particular ecological niche for these defense systems. The Cas13b proteins within these systems had a pairwise sequence identity of only 34.6%. However, they appeared to cluster together within a monophyletic group (**Figure 3A**). This group includes certain Cas13b effectors not associated with RTs, such as those found in *Prevotella* sp. MA2016 and previously described as a variant of the subtype VI-B1 system (Smargon et al., 2017). The Cas13b effector proteins within this branch are found predominantly in bacteria from phylum Bacteroidetes. Furthermore, the linked RTs are phylogenetically more closely related to those associated with the adaptive machinery of type III systems in RT clade 2, as detailed by Toro et al. (2019a). These systems are commonly found in genus *Bacteroides*, suggesting a potential source for their recruitment.

**Figure 3.**
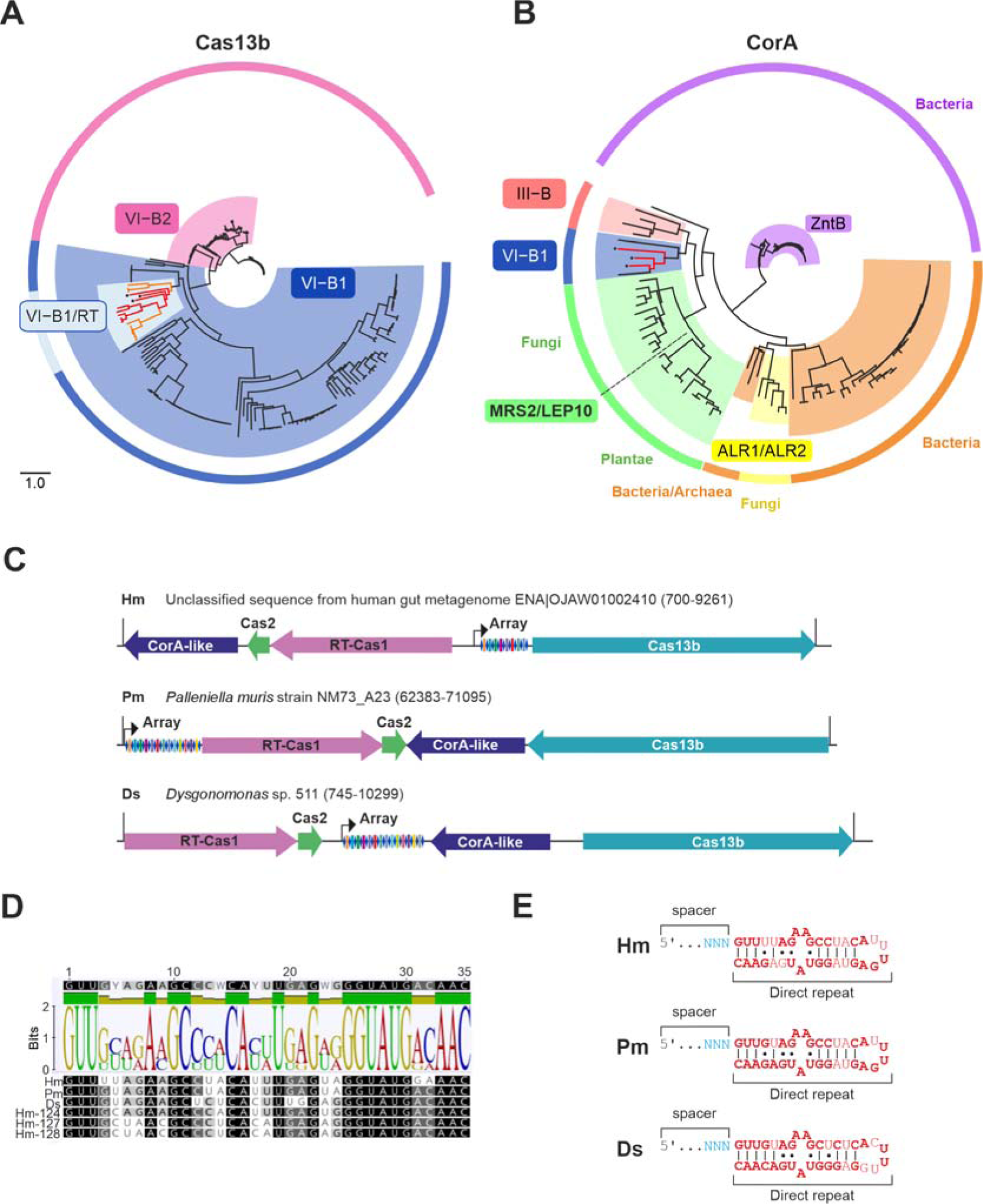
CRISPR-Cas type VI-B systems associated with RTs. **(A)** Schematic phylogenetic tree constructed from the alignment of 228 Cas13b protein sequence variants, distinguishing between subtypes VI-B1 (loci with Csx27) and VI-B2 (loci with Csx28) represented by the blue and pink backgrounds, respectively. A distinctive cluster within subtype VI-B1, characterized by the absence of Csx27 loci (light blue background), appeared to be the origin of the variant associated with RTs (red branches). Black dots indicate the *cas13b* genes within the loci studied here. The corresponding tree newick file is provided as **File S1**. **(B)** Schematic phylogenetic tree derived from the alignment of 124 sequences, including the magnesium transport protein CorA (depicted in dark blue), its related protein, the zinc transport protein ZntB (in purple), the mitochondrial inner membrane magnesium transporter Mrs2 (in green), and the magnesium transporters Alr1 and Alr2 (in orange) from the IPR002523 protein family. Branches corresponding to CorA-like proteins associated with CRISPR-Cas III-B systems are highlighted in coral, whereas those linked to subtype VI-B systems are indicated in blue. Black dots indicate the *CorA*-like genes within the loci studied. The corresponding tree newick file is provided as **File S2**. **(C)** Genomic arrangement of the three type VI-B-RT systems selected from a metagenomic assembly (Hm, ENA|OJAW01002410|OJAW01002410.1), *Palleniella muris* NM73_A23 (Pm, SRZC01000020), and *Dysgonomonas* sp. S11 (Ds, NZ_QVMQ01000048) studied here. The corresponding gene locus tag (final coordinates) is indicated. The black arrows indicate the orientation of the predicted array promoters. Genes are drawn to a proportional scale. **(D)** Alignment of the sequences of the direct repeats (RNA form) of the six arrays associated with RT-Cas1 (ordered as described in **Table S1**) including the resulting Logo graphic. **(E)** Predicted secondary structure of the crRNA substrates of the CRISPR arrays of the three systems studied (in bold, conserved positions). Direct repeat nucleotides are colored in red and spacer nucleotides are shown in light blue (the full spacer is not shown). Watson-Crick base-pairing is denoted by black lines; non-Watson-Crick base-pairing is denoted by dots. This structure is consistent with those previously described for canonical type VI-B arrays (Smargon *et al*. 2017 and Slaymaker *et al*. 2019).

The two known variants of subtype VI-B systems, VI-B1 and VI-B2, can be differentiated on the basis of the presence of additional small accessory proteins containing predicted transmembrane (TM) helices, known as Csx27 (VI-B1) and Csx28 (VI-B2). These accessory proteins regulate Cas13b-mediated RNA interference (Smargon et al. in 2017; VanderWal et al. 2023). These variants are believed to have diverged during the course of evolution, leading to different architectural features associated with these unique accessory transmembrane proteins (Shmakov et al., 2017). Interestingly, the type VI-B systems within the branch, including those associated with RTs, are characterized by the absence of Csx27 and Csx28 accessory proteins. Instead, the Cas13b effector protein appears to be linked to upstream WYL domain-containing proteins or, in the case of *Prevotella* sp. MA2016 and *Alitispes* sp. isolate RGIG9019, a *corA*-like gene downstream from the encoded Cas13b effector, which serves as a distinctive feature. Not only do the type VI-B systems associated with RTs include the CRISPR array and the adaptation module, but they also appear to be linked to CorA-encoding loci with diverse genomic architectures. These CorA proteins are characterized by two or three transmembrane helices at their C-terminal ends and they have an origin in common with those encoded by genes identified at various subtype III-B loci, as previously reported (Shmakov et al. in 2018, Chi et al. in 2023) (**Figure 3B**). There may therefore be a functional association between CorA and these type VI-B/RT systems, linking their defense mechanisms to membrane processes.

### Spacer acquisition in CRISPR type VI-B systems with adaptation modules including RT-Cas1 fusion proteins

Six of the eight VI-B/RT-Cas1 systems identified here comprises unique CRISPR arrays, each containing two to 13 spacers of 31-37 nucleotides in length (**Table S**1). The direct repeats in these CRISPR arrays present characteristic features unique to Cas13b-associated CRISPR arrays (**Figure 3D** and **3E**). They are conserved in size, sequence, and structure, with a length of 35 nt, complementary sequences (GUUG and CAAC) at the ends of the repeat, and a U-rich stretch in the open-loop region, resulting in a predicted secondary structure mediated by intramolecular base-pairing and resembling that of Bz- and Pb-*cas13b*-associated CRISPR arrays (Smargon et al. 2017 and Slaymaker 2019). These data suggest that VI-B/RT-Cas1 CRISPR arrays are probably associated with and processed by their corresponding Cas13b effector proteins (see below).

Three of the eight type VI-B CRISPR-associated RT systems identified here appeared to be complete (**Figure 3C**; **Table S1**). We assessed the ability of these systems to acquire new spacers by inserting the genes encoding their RT–Cas1 and Cas2 proteins into a constitutive expression plasmid (pUC) along with a stretch of at least 250 nt of the corresponding predicted CRISPR array containing only one repeat and the next spacer. Using the spacer acquisition assay previously described for type VI-A systems, we found that the system corresponding to *Palleniella muris* (Pm) was the only one of the three orthologous systems that actively acquired new spacers, with an efficiency greater than that of the CH-type VI-A/RT-Cas1-associated system. The other two types of VI-B/RT-Cas1 systems had much lower levels of activity (more than two magnitudes lower for the *Dysgonomonas* sp. construct) or no activity at all (for the human gut metagenome assembly-derived construct) (**Figure 4A**). Constructs lacking the orthologous *cas2* gene, or with a YAAA mutation in the catalytic active site of the RT (RT^-^) or an E655A mutation in the Cas1 active site had more than 90% lower rates of spacer acquisition (**Figure 4B**). *In vitro* RT assays revealed that the RT^-^ mutant had only background levels of RT activity, whereas the RT activity of the Cas1 mutant was only slightly lower than that of the wild-type protein (**Figure 4C**). Taken together, these findings suggest that the acquisition of new spacers by the Pm type VI-B system with an associated RT is independent of other CRISPR adaptation units present in the *E. coli* genome.

**Figure 4.**
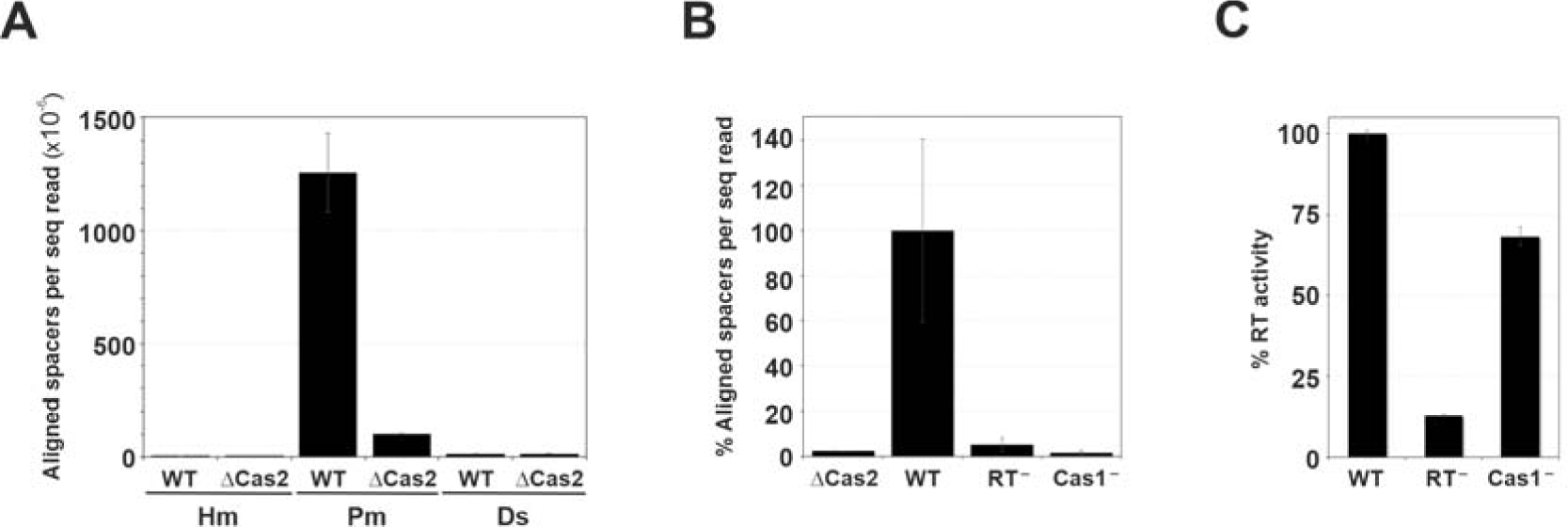
Spacer acquisition in type VI-B CRISPR systems. **(A)** Spacer acquisition by the adaptation modules of the three type VI-B systems (described in Figure 3A) in the heterologous *E. coli* system. Black bars represent the frequency of unique aligned spacers per million reads for each RT-CRISPR-Cas system: (Hm) metagenomic assembly (Pm) *Palleniella muris*, and (Ds) *Dysgonomonas* sp. A negative control is included for each system in which the *cas2* gene is deleted (ΔCas2). The bars indicate the range for at least three biological replicates. **(B)** Percentage of aligned spacers relative to the *P. muris* wild-type (WT) system normalized per sequencing read of the different mutants is represented: ΔCas2 lacks the Cas2 protein in the adaptive operon; RT-, RT-Cas1 protein with a mutation affecting the catalytic core for RT activity; YADD → YAAA; and Cas1-, the RT-Cas1 coding gene is modified to replace a glutamate residue with alanine (E665A), a substitution predicted to abolish Cas1 function. **(C)** Exogenous RT activity measured as in Figure 1. Polymerization is presented as a percentage of the RT-Cas1 activity of the WT protein (WT). RT-: mutation affecting the catalytic core for RT activity and Cas1-, mutation affecting cas1 endonuclease activity. The data were derived from three technical replicates and at least one biological replicate and are presented as the mean ± the standard error of the mean (SEM).

### Features of spacer acquisition by type VI-B/RT CRISPR-Cas systems

We further investigated the characteristics of spacer acquisition by the Pm type VI-B/RT-Cas1 system, by analyzing the pool of newly acquired spacers. Over 95% of the spacers were between 32 and 35 base pairs (bp) long, consistent with the length distribution of the natural spacers present in these arrays (**Figure 5A**). In addition, their median GC content was consistent with that of the two templates used: the *E. coli* genome (**Figure 5B**) or the plasmid (not shown). The acquisition process showed no bias towards rRNA genes and was not correlated with the level of gene expression (not shown). Moreover, there was no bias towards a specific antisense orientation of coding sequences, indicating an absence of complementarity between the newly acquired spacers and the predicted messenger RNA (**Figure 5C**). A significant deviation of the expected GC content was observed at various positions within the spacer unit (**Figure 5D**). A symmetric bias therefore emerged, with a stretch of two to three nucleotides rich in AT residues observed at both ends of the spacers. A similar symmetric bias of AT positions was observed at the +17 position of the spacer, not only for spacers originating from the genome, but also for those originating from the plasmid. Interestingly, the first nucleotide flanking the 5’- and 3’-ends of the acquired spacers in the protospacer displayed a preference for G and C nucleotide residues, respectively.

**Figure 5.**
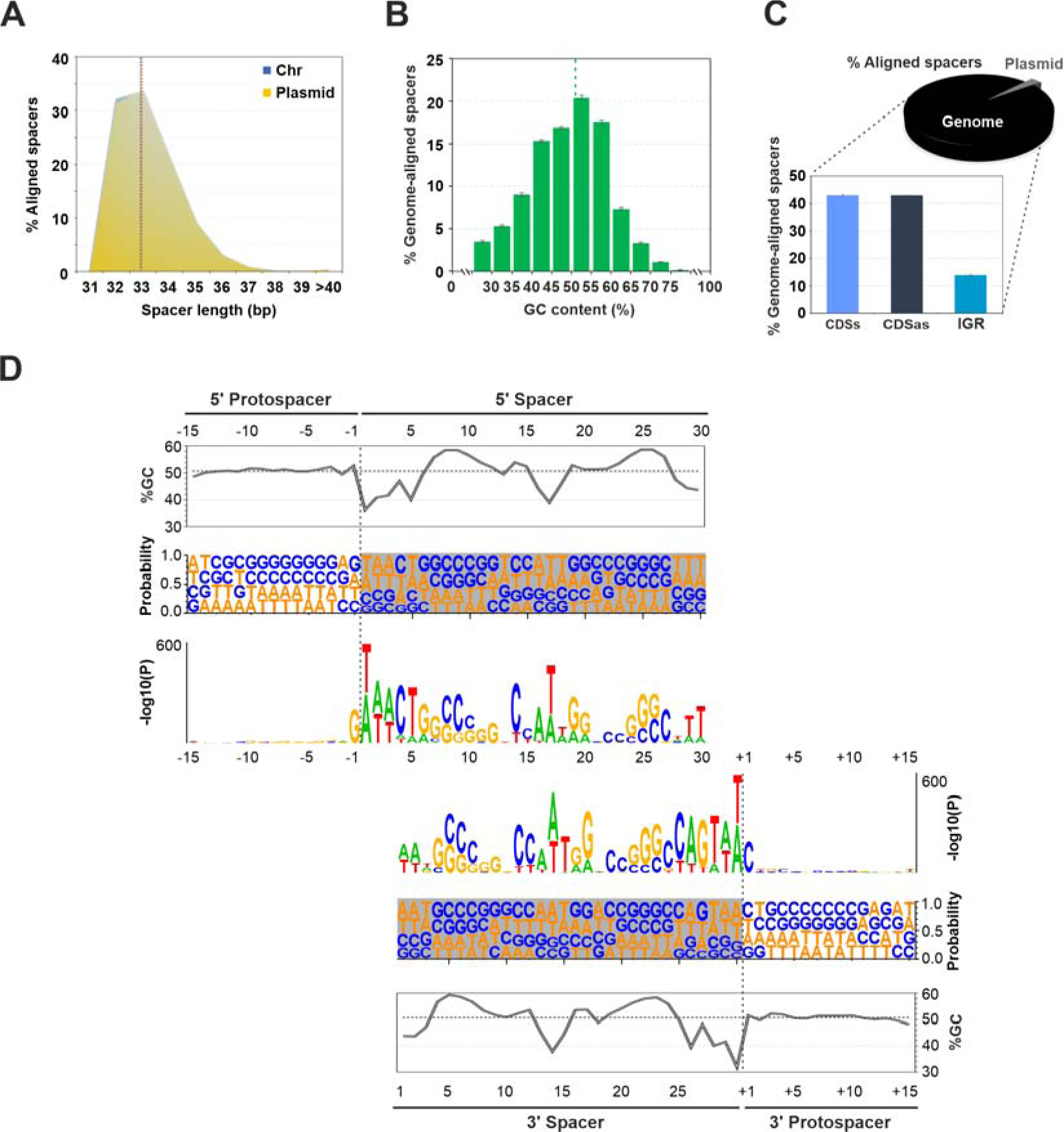
Characteristics of the spacers acquired by the *Palleniella muris* adaptation system. **(A)** Distribution of newly detected spacers by length. Blue traces match reads identified in the *E. coli* chromosome and orange background correspond to those matching the sequence encoding the adaptation operon in the plasmid. **(B)** GC content distribution in the genome-aligned spacers. The dotted lines represent the mean GC content of the genome. **(C)** Strand bias in pools of newly acquired spacers relative to the source transcript. Bars plot represents the proportion of newly acquired spacers with the wild-type RT-Cas1 in the sense or antisense strand of coding genes or in intergenic regions of the *E. coli* genome (*n* = 9). Above, a pie chart with the proportion of spacers aligning with the genome and the plasmid. **(D)** Protospacer composition analysis for GC content, frequency and statistical probability. The spacer sequence is shaded and the spacer boundaries are marked as dotted lines. Due to the different lengths of the detected spacers, the protospacer sequence is split into a 5’ protospacer (upper panel) and a 3’ protospacer (lower panels). The dotted line on the GC content plot corresponds to the mean % GC in the *E. coli* genome. On the statistical probability graph [-log_10_ (P)], nucleotides are represented such that their size indicates their relative probability (average across all positions) using k-mer length=1, 2, 3, 4 and a binomial test. Data represent spacers merged across *n* = 9 biological samples based on <75000 data.

We investigated the possible impact of the associated Cas13b effector protein on spacer acquisition patterns by generating novel plasmid constructs (**Figure 6A**). We inserted the RT-Cas1/Cas2 adaptation locus into an arabinose-inducible plasmid backbone (the pBAD plasmid series), immediately downstream from the *cas13b* gene. The Cas13b protein of *P. muris* consists of 1265 amino acids, with two predicted HEPN sites responsible for predicted collateral RNase activity identified as ‘RNXXXH’ motifs (positions 148-153 and 1214-1219). Furthermore, structural alignment with the PbuCas13b protein revealed the presence of a potential Lid region, a domain that caps the 3’ end of the crRNA with two beta hairpins. Within this region, the Pm Cas13b Lys (K) 444 is predicted to correspond to K393 in PbuCas13b, a critical residue of the Lid domain responsible for crRNA processing (Slaymaker et al., 2019).

**Figure 6.**
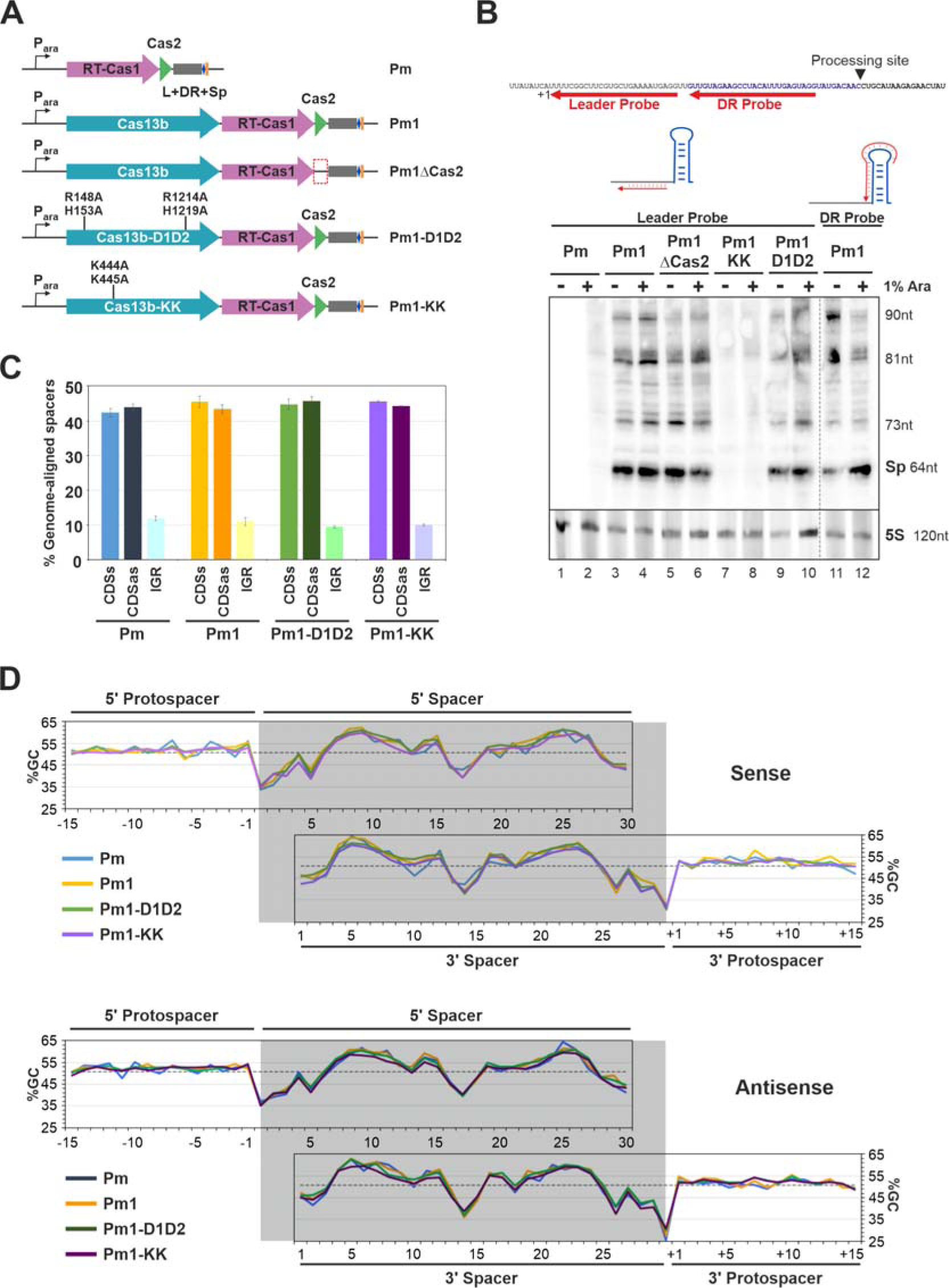
Contribution of Cas13b to spacer acquisition in *Palleniella muris*. **(A)** Schematic diagram of the pBAD derivatives containing Cas13b sequences. (Pm) A plasmid containing the RTCas1-Cas2 sequences and a minimal unit of the CRISPR array expressed under the control of an arabinose-inducible promoter. (Pm1) Pm plasmid with the gene encoding the corresponding Cas13b (Pm1), Cas13b mutated at the two HEPN sites (Pm1-D1D2) and Cas13b mutated at the two K residues of the Lid region (Pm1-KK). An additional control in which the *cas2* gene was removed (Pm1ΔCas2) was included. **(B)** Cas13b encoded by the *P. muris* VI-B locus processes the CRISPR array. Northern-blot analysis of exponential growth phase *E. coli* cell cultures with and without 1% arabinose, with the constructs depicted in (A). Two different antisense strand-specific oligonucleotides (leader probe, lanes 1-10; direct repeat probe, lanes 11-12) were used to probe total RNA. The figure shows all the hybridization signals detected with each probe. The expected sizes of the bands are indicated. At the bottom, a 5S RNA from the different samples is included as a control for RNA loading. **(C)** Strand bias in pools of newly acquired spacers relative to the source transcripts. Proportion of newly acquired spacers with the four different constructs in the sense (CDSs) or antisense (CDSas) strand of coding genes or in intergenic regions (IGR) of the *E. coli* genome (*n* = 3). **(D)** Protospacer composition analysis for GC content at each position along the acquired spacers in the sense (above) or antisense (below) strand of coding genes obtained from *E. coli* cultures with the various constructs (figure legend) is represented. The spacer sequence is shaded. Due to the different lengths of the detected spacers, the protospacer sequence is split in a 5’ protospacer (upper panel) and a 3’ protospacer (lower panels). Data represent spacers merged across *n* = 3 biological samples. The dotted line on the GC content plot represents the mean % GC for the *E. coli* genome.

Before analyzing the effect of Cas13b on acquisition, we checked that the pre-crRNA was processed (**Figure 6B**). A northern-blot analysis of total RNA from a plasmid containing the *cas13b* gene yielded bands corresponding to the mature forms of crRNA, demonstrating the occurrence of processing. This processing was abolished in the K444A-K445A mutant but maintained in the D1-D2 HEPN site double mutant, as expected. Furthermore, the expression of the Cas3b effector in the presence of the associated CRISPR array leads to cell dormancy or death, as demonstrated by growth suppression, probably reflecting the collateral RNase activity (Abudayyeh et al., 2016; East-Seletsky et al., 2016) caused by the interference complexes (**Figure S2**). These findings indicate that the PmCas13b effector protein is functional in the *E. coli* host. We then used these constructs to analyze the pattern of spacer acquisition. The presence of the Cas13b effector gene did not alter the spacer acquisition profile, and we observed no bias towards targeting the complementary strand relative to the gene-coding sequences or intergenic regions (IGRs) (**Figure 6C**). Moreover, the nucleotide position data were similar to those obtained in the absence of the effector locus (**Figure 6D**). These findings suggest that, unlike its counterpart in the type II-C system, the type VI-B effector protein is not involved in the adaptation process.

### The *P. muris* type VI-B/RT-Cas1 CRISPR adaptation module acquires spacers directly from RNA

Only three type III CRISPR-associated RT systems have been shown to acquire spacers directly from RNA *in vivo*: a III-A system from *Marinomonas mediterranea* (Mm), and two III-D systems, one from *Fusicanibacter saccharivorans* (Fs) and the other from *Vibrio vulnificus* (Vv) (Silas et al., 2016; Schmidt et al., 2018; González-Delgado et al., 2019). We investigated whether the CRISPR adaptation module of a novel RT-Cas1 system, the Pm type VI-B/RT-Cas1 system, could acquire spacers from RNA molecules. We used a previously described acquisition assay, in which we searched for spacers harboring the ligated exon junction of the self-splicing *td* group I intron, a ribozyme that catalyzes its own excision from the original transcript and is not present in DNA (Silas et al., 2016; González-Delgado et al., 2019). We designed a construct containing the *td* intron immediately upstream from the adaptation module operon, expressed under a constitutive promoter (**Figure 7**). As in previous designs, the detection of the exon junction as a DNA spacer was considered to provide evidence that the system was indeed able to acquire spacers directly from RNA. We then searched for newly integrated spacers and identified more than 50,000 new spacers mapping onto plasmids or to the *E. coli* genome. We discovered two unique spacers spanning the splice junction (**Figure 7**), confirming, as observed with type III CRISPR-associated RT systems, that type VI-B/RT-Cas1 CRISPR-Cas adaptation modules can indeed acquire spacers from RNA molecules in *E. coli*.

**Figure 7.**
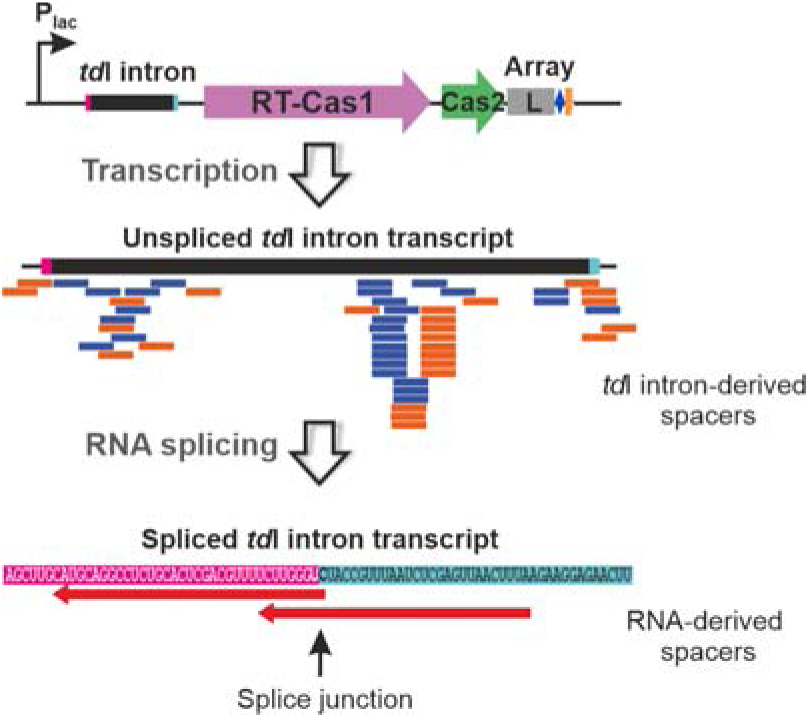
Spacer acquisition from RNA by the *Palleniella muris* adaptation system. Above, schematic diagram of *td* intron-containing constructs. We determined whether the spacers originated from RNA, using a self-splicing transcript that produces an RNA sequence junction not encoded by DNA. High-level coverage of spacers annealed with the *td*I gene is shown (blue and orange bars correspond to sense and antisense spacers, respectively). At the bottom, alignment of newly acquired exon junction-spanning spacers (considered to have been acquired from an RNA target), indicated as red arrows. Note that both spacers are in the antisense orientation relative to the direction of transcription of the *td*I intron. Flanking exons are shaded in pink (5’) and turquoise (3’).

## DISCUSSION

We have previously described CRISPR-Cas type VI-A systems associated with RT-Cas1 fusion proteins (Toro et al., 2019b). Here, we report the presence of a subtype VI-B1 variant of the CRISPR-Cas system associated with RT-Cas1 fusion proteins in gut metagenomes and genome assemblies. In addition to the Cas13b effector protein, this CRISPR-Cas variant carries a characteristic type VI CRISPR array, an RT-Cas1/Cas2 integrase complex, and a linked CorA-encoding locus. We investigated the acquisition of spacers *in vivo* by the RT-Cas1/Cas2 integrase complex machinery associated with type VI-A and VI-B systems. Our findings indicate that these variant type VI systems are fully functional in terms of the acquisition of new spacers from DNA. Furthermore, we demonstrated that the Pm type VI-B system associated with an RT-Cas1 fusion protein can acquire RNA-derived spacers in an RT-dependent manner. The associated Cas13b effector can process precursor crRNA, but does not seem to be involved in the acquisition process. Spacer acquisition by both type VI-A and VI-B RT-Cas1 systems appears to display a preferential targeting of 32 to 35 bp sequences with AT-rich sequences at both ends. The type VI-B systems also appear to prefer the protospacer to be flanked by G and C residues at the 5’and 3’ends, respectively. We observed no particular acquisition bias for highly transcribed loci or genomic regions.

All the type VI-B1/RT systems reported here were derived from gut metagenomes. Preliminary analyses of the presence of this type VI-B1/RT system variant in specific gut metagenomes, such as the broad early-life human gut microbiome (Zeng et al., 2022), have suggested that it accounts for only a small proportion (4.54%) of canonical type VI-B1 systems. This variant was found in only two of the 6122 fecal metagenomes sampled (not shown). These findings raise questions about the significance of type VI-B1/RT systems in this particular habitat and whether their presence is related to specific threats to human health or environmental factors, diet or lifestyles. Further studies are required to address this question.

The phylogenetic relationships between the VI-B1-associated RTs and the CorA-encoded loci, and their taxonomic distribution predominantly in Bacteroidetes suggest that the adaptation module (RT-Cas1, Cas2) and CorA of type VI-B systems are probably derived from those found in type III systems. This evolutionary transition may have occurred in response to environmental pressures within specific habitats, such as the intestinal tract of animals. A plausible scenario involves a single evolutionary event with gains and losses. According to this scenario, the closely related type VI-B1 systems from *Prevotella* sp. MA2016 and *Alitispes* sp. isolate RGIG9019, both containing a *corA*-like gene downstream from the gene encoding the Cas13b effector protein, have lost the RT-Cas1/Cas2 adaptation module. The acquisition of CorA proteins, characterized by two or three transmembrane helices, probably rendered the presence of the Csx27 loci typical of type VI-B1 systems redundant, eventually leading to their loss. However, other explanations for these evolutionary dynamics remain plausible.

It has been suggested that the CorA effector associated with type III-B systems may bind to the generated SAM-AMP, leading to cell dormancy or death (Chi et al., 2023). On the other hand, it has been suggested that the VI-B1-associated ancillary Csx27 protein, which downregulates the activity of Cas13b, may form membrane channels for ssDNA uptake, and that its nascent transcripts are degraded by the VI-B1 interference machinery (Makarova et al., 2019). Additionally, the Csx28 protein associated with VI-B2 systems forms pores, and its antiviral activity requires sequence-specific cleavage of viral mRNAs by Cas13b leading to membrane depolarization (VanderWal et al., 2023). The presumed role of CorA in the interference step in type VI-B/RT-associated systems needs to be determined, but we are more inclined towards a role similar to that shown by Csx28.

It remains challenging to understand spacer acquisition in type VI systems, due particularly to the absence of an adaptation module in many of these systems. Crosstalk between CRISPR-Cas systems and the *trans*-adaptation model is a concept that could shed some light on the mechanisms operating in some type VI systems lacking adaptation genes. For instance, studies on *Flavobacterium columnae* have shown that, despite the presence of an acquisition-deficient subtype VI-B system, this microorganism can still acquire spacers in *trans*, possibly through interactions with other CRISPR-Cas systems (Hoikkala et al., 2021). However, additional experimental investigations of adaptation in type VI systems are required before any firm conclusions can be drawn. The identification of type VI-A- and type VI-B-associated RT-Cas1/Cas2 systems enabled us to explore how type VI systems acquire spacers *in vivo* and whether they can acquire spacers from RNA sources. Our findings suggest that these variants of the type VI system are fully functional for the acquisition of new spacers from DNA. Interestingly, this acquisition does not appear to involve the adaptation modules of any other CRISPR-Cas systems present. Spacer acquisition is instead mediated by the associated RT-Cas1/Cas2 integrase complex. Furthermore, unlike type II-C systems (Rousseau et al., 2018), our results indicate that the acquisition process of these systems does not require the associated effector proteins (Cas13a or Cas13b).

Spacer acquisition was inefficient in type VI-A/RT systems, but acquisition efficiency was higher in the Pm type VI-B/RT system, making it possible to characterize the spacer acquisition process *in vivo* with greater confidence. This particular system is also predicted to be fully functional for the interference step, as demonstrated by the ability of its Cas13b effector to process the associated pre-crRNA, leading to a suppression of cell growth and suggesting than an interference complex may be formed. Interestingly, acquisition by the Pm system is not biased towards highly transcribed regions or specific genomic locations, indicating an absence of preference for spacers potentially targeting mRNA. This observation may reflect a preference for dsDNA over RNA substrates in this type of VI-B/RT-associated system, as shown *in vitro* for the *Thiomicrospora* type III Cas6-RT-Cas1-Cas2 complex (Wang et al., 2021). In both types of VI-A and VI-B/RT-associated systems, the ends of the spacers tended to be enriched in AT nucleotides. Furthermore, in the Pm-type VI-B/RT system, the target sequences displayed a preference for G and C nucleotides at the −1 and +1 positions, respectively, of the protospacer. This suggests that the integrase complex may make use of a two-sided protospacer flanking sequence (PFS) for target recognition. However, the significance of this recognition for the interference step remains to be determined. This system was nevertheless able to acquire spacers from RNA molecules, consistent with expectations for CRISPR-associated RT systems. It is difficult to assess the efficiency of this reaction *in vivo*, as it may depend on the host and the availability of spacer sources. It is therefore not currently possible to draw any firm conclusions about the primary source of spacers in these systems.

Type VI effector proteins cleave RNA targets only (Abudayyeh et al., 2016; Smargon et al., 2017; Shmakov et al., 2015; Shmakov et al., 2017). Many of these systems have no adaptation module, but the presence of such a module in others suggests a possible ability to acquire spacers directly from RNA phages. The discovery of novel type VI-B systems associated with RT-Cas1 fusion proteins made it possible to decipher the details of spacer acquisition in type VI systems and to show that they are indeed able to acquire spacers from RNA sources. CRISPR-Cas systems are a component of microbial defense mechanisms. The presence of type VI-B/RT systems in the gut microbiome may, therefore, indicate interactions with RNA viruses or other novel RNA genetic entities (Zheludev et al., 2024). By studying the association between these CRISPR-Cas systems and specific health outcomes or disease states, we may be able to identify novel biomarkers or indicators of gut health and disease susceptibility.

## Acknowledgments

This work was supported by Grant PID2020-113207GB-I00 funded by MCIN / AEI / 10.13039/501100011033 and Grant P20_00047 funded by Consejería de Transformación Económica, Industria, Conocimiento y Universidades (Junta de Andalucía) with FEDER founds. We thank Ascensión Martos Tejera for providing technical assistance. We also thank to Fernández-González, A.J. for developing the custom scripts used in this work.

## Authorścontributions

MDM, FMA and NTG designed the experiments and contributed to the writing-original draft, its review, and editing, with contributions from MRM. MDM and VMC performed the experimental work on type VI-A and type VI-B systems, respectively. MRM performed database analysis to identify additional type VI systems associated with RT. NTG identified type VI-B systems associated with RT-Cas1 fusion proteins and performed the phylogenetic analyses. MDM, FMA and NTG contributed to the interpretation of experimental results.

## Declaration of interest

The authors have no competing financial interests to declare.

## MATERIALS AND METHODS

### Strain and culture conditions

The *E. coli* strains used in this study were DH5α (Bethesda Research Laboratories) for cloning and Rosetta 2 (DE3) (Novagen) for protein production. For spacer acquisition assays, several different *E. coli* strains were used, as appropriate for the experiment: HFR3000 F+ or HMS174 (DE3) for pUC-based experiments and BW25113 for pBAD-based experiments to ensure effective arabinose induction. The bacteria were grown in LB medium (10 g/l tryptone, 5 g/l yeast extract, and 5 g/l NaCl) at 37°C with orbital shaking. Antibiotics were added as required (ampicillin, 200 µg/ml; chloramphenicol, 50 µg/ml; and kanamycin, 50 µg/ml).

### Plasmid construction for spacer acquisition assays and protein production

The plasmids used for spacer acquisition experiments were derived from the pBAD or pUC backbone, depending on whether the construct needed to include the coding sequence for the effector protein Cas13. Rk plasmids were obtained by cloning PCR products obtained with the indicated primers and genomic DNA supplied by the German Collection of Microorganisms and Cell Cultures GmbH, DSMZ (DSM#19783) as a template. The remaining systems were generated by and obtained from GenScript Biotech Corporation (EU Headquarters). Unlike the Rk system, the entire RT-Cas1_Cas2 adaptation module (wild-type and mutant) was codon-optimized for expression in *Escherichia coli* under the control of a constitutive promoter in the pUC backbone. We ordered each individual protein cloned as an SphI/SalI block on a scaffold digested with SphI/XhoI. In each case, the CRISPR array consisted of the leader region, the first DR and the first spacer (**Figure S1A**). The Cas13-encoding gene was ordered directly in an inducible pBAD backbone for induction of the expression of the complete CRISPR-Cas operon.

The plasmids used for the production and purification of proteins were based on the pMal-Flag-3C backbone (González-Delgado et al., 2019). RT-Cas1 was amplified from the corresponding pUC vector (or genomic DNA from Rk) and inserted into pMAL-3C-Flag as a BamHI/BglII fragment. The *R. kholense* RT-deficient mutant was obtained by two rounds of PCR with oligonucleotide primers containing the desired nucleotide substitution. The plasmids and oligonucleotides used in this study are listed in **Tables S2-4**, and the sequences of the different constructs used are also included in **File S3**. The *td* intron construct was obtained by inserting the 393 bp intron sequence and its native exons into pUC_Pm immediately upstream from the RT-Cas1 coding sequence. The receptor plasmid was digested with XhoI and dephosphorylated. The sequence containing *td*I was inserted as a SalI/XhoI fragment from pG-tdI (Δ1-3)s (Martínez-Abarca et al., 2004).

### Protein purification

We transformed the *E. coli* strain Rosetta2 (DE3) with various pMal-Flag derivatives carrying RT-Cas1-coding genes. Single colonies were cultured overnight at 37°C in LB medium supplemented with ampicillin, chloramphenicol, and 0.2% glucose, with shaking. A 1/100 dilution of the resulting culture was used to inoculate a flask containing 50 ml LB supplemented as indicated, which was then incubated at 37°C, with shaking, until the culture reached the exponential growth phase. The cultures were then induced by adding IPTG to a final concentration of 1 mM IPTG and incubating overnight at 20°C. Cells were harvested by centrifugation, the pellet was washed with PBS buffer (137 mM sodium chloride, 2.7 mM potassium chloride, 10 mM sodium phosphate dibasic, 1.8 mM potassium phosphate monobasic), and quickly frozen in liquid nitrogen. Cells were lysed in column buffer (CB: 20 mM Tris–HCl (pH 7.5), 200 mM NaCl, 1 mM EDTA, 1 mM DTT and 1x EDTA-free protease inhibitor (Roche)) with a French press. The lysate was cleared by centrifugation (16,000 × *g* for 15 min at 4°C). Proteins were purified by affinity liquid chromatography. Amylose resin was placed in a 10 mL Econo-Pac disposable column (BioRad) and washed several times with CB. Crude protein extract was incubated with the column for 2h at 4°C with horizontal shaking. The bound proteins were eluted by incubation in CB supplemented with 10 mM maltose for 1h at 4°C. The proteins were concentrated with an Amicon ultracentrifugation filter (YM-30) and dialyzed against storage buffer (SB: 10 mMTris–HCl (pH 7.5), 1 mM DTT, 50% glycerol). Protein extracts were stored at −80°C until required. Protein concentration was determined by the Bradford method (Bio-Rad), according to the kit manufacturer’s protocol.

### Exogenous RT assay

Reverse transcriptase reactions were performed as previously described (Muñoz-Adelantado et al., 2003). Briefly, 500 ng of protein extract was incubated with poly(rA)/oligo(dT)_18_ or poly(rA) substrates in the presence of RT buffer (10 mM KCl, 25 mM MgCl_2_, 50 mM Tris–HCl (pH 7.5), 5 mM DTT) and 2.5 µCi [α-^32^P] dTTP (800 Ci/mmol; Perkin Elmer) for 10 min at 37°C. The reaction was stopped by spotting 8 µL of the reaction mixture onto Whatman DE81 paper. The paper was dried, washed with 250 ml 2×SSC, and radioactivity was quantified with a scintillation counter (Liquid Scintillation Analyzer Tri-Carb 1500, Packard).

### Spacer acquisition assay

The appropriate *Escherichia coli* strain was transformed by electroporation with a plasmid expressing the desired adaptive operon, basically the RT-Cas1-Cas2-CRISPR array and its derivatives (**Table S2**). At least nine individual colonies were cultured overnight at 37°C in liquid LB medium supplemented with ampicillin. The culture was diluted 1:500 in LB medium, with the addition of 0.005% arabinose if required for induction, for 14–18 h at 37°C. The bacterial cells were harvested by centrifugation, and the plasmids were isolated by standard procedures. Leader proximal spacers were amplified by PCR with 2 ng plasmid DNA as the template, a forward primer binding to the leader sequence, and a reverse primer binding to the first native spacer (**Table S4**) under the following cycling conditions: 94°C for 2 min; 30 cycles of 94°C for 20 s and 58-62°C for 20 s. The dominant PCR product was an unexpanded array. We therefore performed a blind purification of the expanded CRISPR array consistent with a band approximately 70 nt larger, with the mi-Gel Extraction Kit (Metabion International AG, Germany). The purified samples were then used for a second round of PCR with divergent primers binding to the DR sequence (DR-PCR Mackenzie et al. 2019; **Figure S1**). An amplicon band was detected only if an acquisition event occurred. Quantification was performed with a Qubit fluorometer (Life Technologies), with analysis on a 2100 Bioanalyzer (Agilent Technologies). Libraries were sequenced with Illumina technology at the Genome Sequencing Unit of IPBLN-CSIC (Granada, Spain). The detection of spacers matching the exon junction after *td* intron splicing was maximized by performing experiments with 30 individual colonies as the starting material for the leader-spacer PCR. The samples were then processed according to the standard procedure, as described above.

### Data processing pipeline

FASTQ files were refined with a custom script and SEED 2.1.1 software: size selection, trimming of adaptor sequences, grouping as unique sequences, and clustering of the unique sequences at a threshold of 90% identity to reduce the experimental background due to PCR bias. Spacers were identified with a custom-written Ruby script determining the sequence between the 3’- and 5’- ends of the DR sequence. Mapping onto the plasmid and genome was performed with Bowtie2.0, with two mismatches allowed. For protospacer sequence analyses, we used a new custom-written Ruby script reporting the 15 nucleotides upstream from the first position matching the spacer sequence and the next 30 nucleotides within the spacer. We also obtained the last 30 positions of the spacer and 15 nucleotides at the 3’ end of the protospacer. Logos were generated with both WebLogo 3.0 (https://weblogo.threeplusone.com/; Crooks et al., 2004) and kpLogo (http://kplogo.wi.mit.edu/; Wu and Bartel, 2017) using default parameters.

### Northern-blot hybridization

RNA was extracted from cultures of *E. coli* BW25113 containing pBAD plasmids with or without sequences encoding the wild-type Cas13b or the corresponding mutants, as previously described (Molina-Sanchez et al., 2006). Bacterial cultures at exponential growth phase (6 ml, 0.2 U OD_600_) were left uninduced or were induced by incubation with 1% arabinose for 4 hours at 37°C. We ran 30 µg of DNAse I-digested total RNA samples on 6% denaturing polyacrylamide gels (8 M urea), which were then blotted onto positively charged nylon membranes (Roche, Germany). The RNA bands were fixed to the filter by UV-crosslinking for 2 minutes. Hybridization probes were radioactively labeled as follows: 100 pmol oligonucleotide was incubated with 10 µCi [γ-^32^P]-ATP in the presence of 10 U T4 polynucleotide kinase (NEB) at 37°C for 2 hours. Non-incorporated radiolabeled nucleotides were removed with Sephadex G25 columns (Cytiva, UK). Four probes were used: Leader Probe (5’-CCTCATTTTCAGCACGAAGCCGAAA-3’), complementary Leader Probe (5’-TTTCGGCTTCGTGCTGAAAATGAGG-3’), DR Probe (5’-CTACTCAAATGTAGGCTTCTACAAC-3’) and complementary DR Probe (5’-GTTGTAGAAGCCTACATTTGAGTAG-3’). The membranes were incubated for one hour in phosphate buffer supplemented with 7% SDS, and hybridization was then performed overnight at 42°C with 5 x 10^6^ Ci probe. Successively more stringent washing steps (2xSSC/0.1%SDS; 1xSSC/0.1%SDS; 0.1xSSC/0.1%SDS) were performed, with each step lasting 15 minutes at room temperature. Hybridization signals were obtained by digital imaging of the exposure plate (Fuji), with visualization on a Personal Molecular Imager FX scanner and analysis with Quantity One software (BioRad).

### Assay of E. coli cell viability

A single colony of cells freshly transformed with one of the constructs in a pBAD-derived plasmid was grown overnight at 37°C in LB supplemented with ampicillin and 1% glucose. The culture was then diluted 1/100 dilution and grown to an OD_600_ of 0.4. Serial dilutions of the culture (10 µl per plate) were then deposited on LB plates supplemented with 1% arabinose and incubated overnight at 37°C.

## List of supplementary data

**Figure S1.**
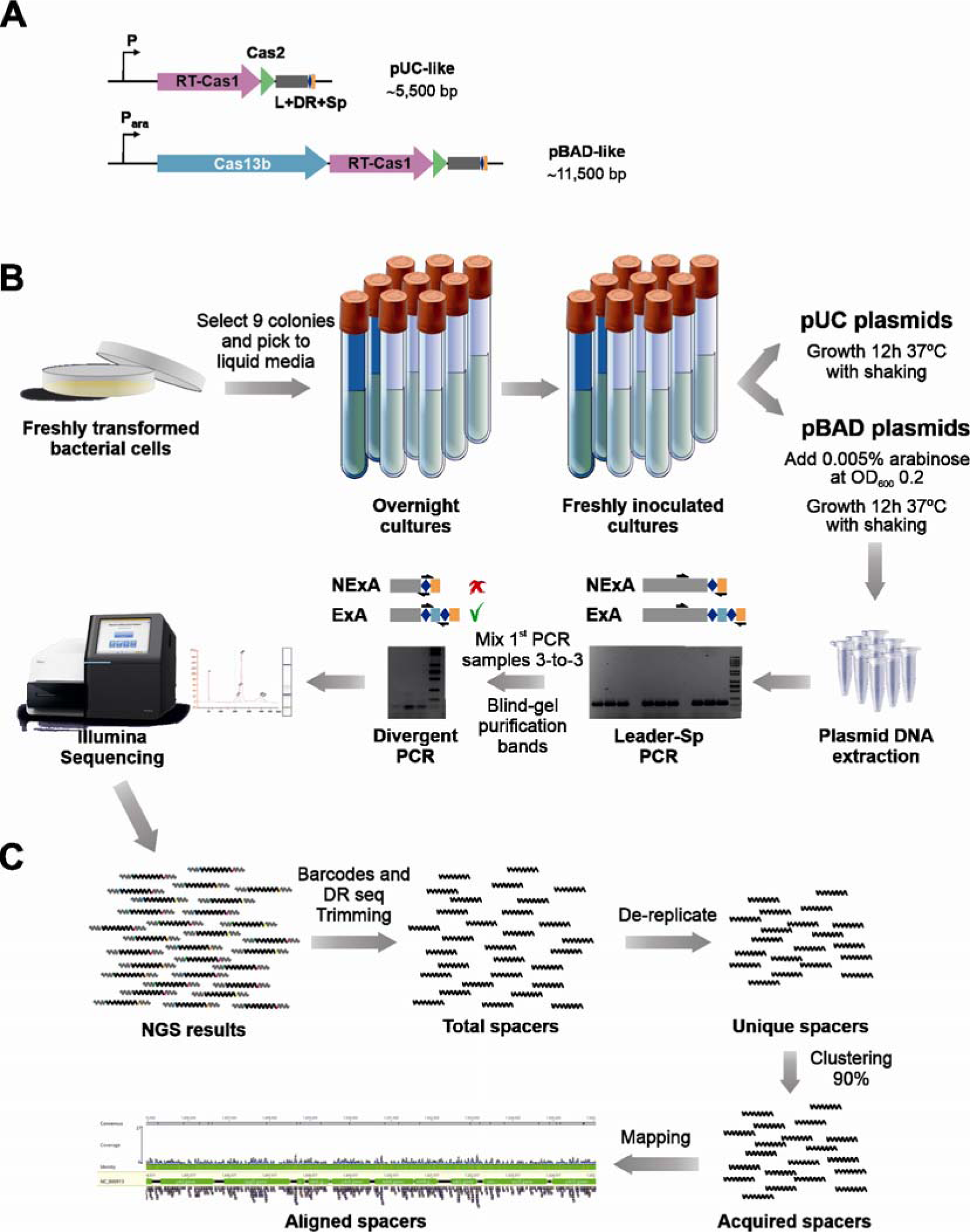
Schematic diagram of the high-throughput spacer acquisition assay. (**A**) Description of the two different examples of adaptation module constructs used in the assays. A pUC-like derivative plasmid with expression under the control of a constitutive promoter; and a pBAD-derived plasmid with expression under the control of an arabinose-inducible promoter. Both types of constructs carry a minimal CRISPR array containing leader–DR–spacer1. (**B**) Experimental workflow (details provided in the methods). Nine single colonies of freshly transformed *E coli* cells were grown overnight in liquid medium at 37°C to late exponential growth phase (for pUC plasmids) or until an OD_600nm_ of 0.2 was reached. We then added 0.005% arabinose and culture was continued for 12 h (for pBAD derivatives). Plasmid DNA was then extracted. These plasmids were the template for a first round of nine individual PCRs with primers binding the leader region (Lfw primer) and spacer 1 (orange block; Sprv primer). These 9 individual PCRs were mixed in groups of three for blind gel-expanded band purification. The purified products were subjected to a second round of PCR (3x) with divergent primers binding to the DR sequence (DR-PCR Mackenzie et al. 2019), oriented such that a product was amplified only if two repeats were present, that is, if the array was expanded. Every amplicon was size-purified and barcoded deep sequencing was performed on the Illumina system. (**C**) NGS sequence processing. FASTQ files for the NGS results were refined with a homemade script and the SEED 2.1.1 software: size selection, trimming of adaptor sequences, removing replicated sequences (unique spacers), and clustering at the 90% identity threshold (acquired spacers). These sequences were mapped onto the plasmid and the genome with Bowtie2.0, with two mismatches allowed, to identify the aligned acquired spacers.

**Figure S2.**
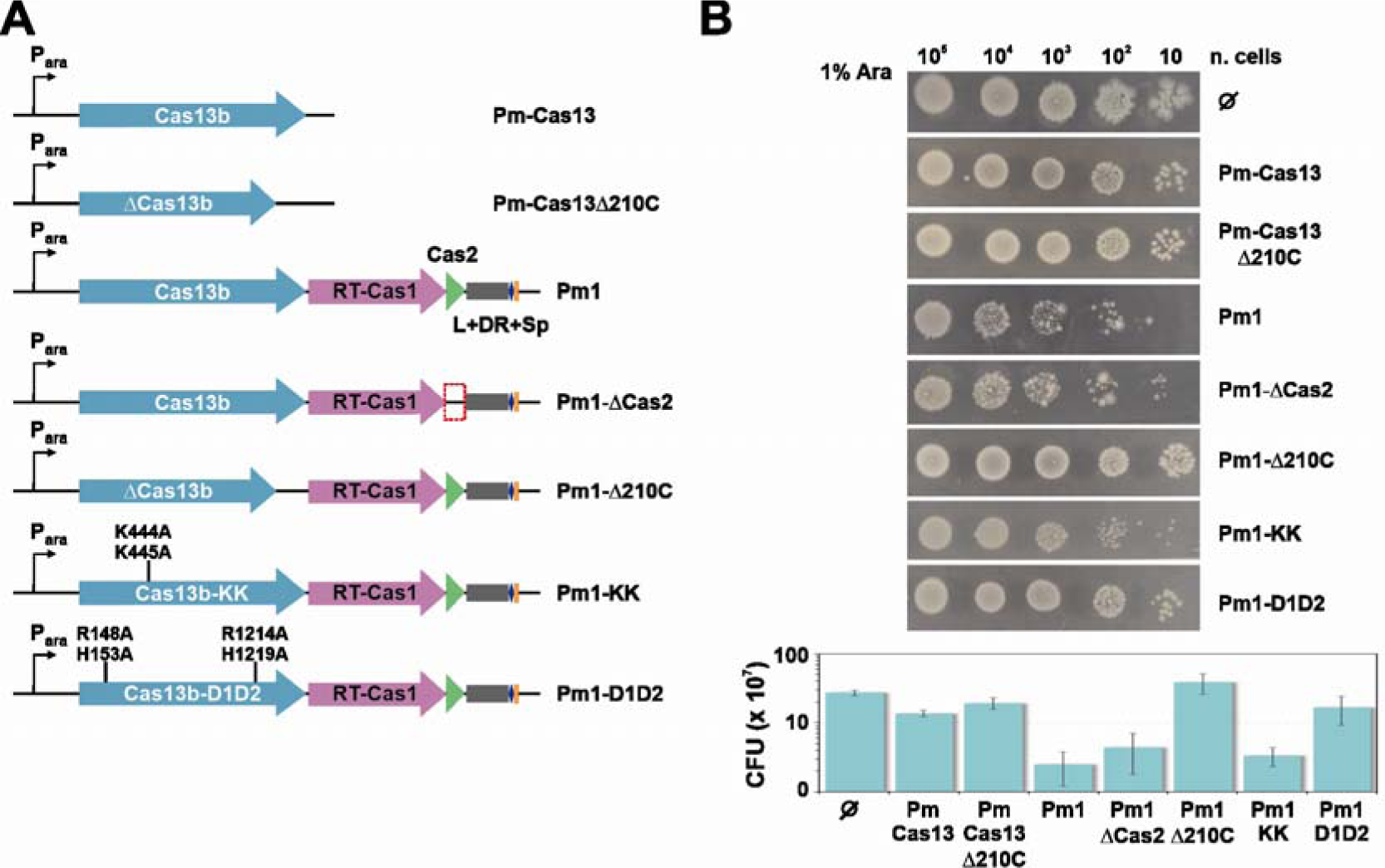
Assay of the viability of *E coli* cells harboring Cas13b-containing constructs. **(A)** Schematic diagram of the pBAD-derived Cas13b-containing constructs used. pBAD plasmids containing only the Cas13b gene under the control of an arabinose-inducible promoter (Pm-Cas13b, wt and Pm-Cas13Δ210C with a frameshift in the *cas13* gene causing a deletion of the 210 C-terminal aa). pBAD plasmids containing the corresponding Cas13b wt gene, the complete adaptation module (RTCas1-Cas2) and a minimal unit of the CRISPR array: (Pm1) WT Cas13b, (Pm1-Δ210C) with frameshift in the *cas13* gene causing a deletion of the 210 C-terminal aa, (Pm1-KK) Cas13b with mutations of the two K residues of the Lid region and (Pm1-D1D2) Cas13b with mutations of the two HEPN sites. An additional control empty vector is included, and a control with deletion of the *cas2* gene (Pm1-ΔCas2). **(B)** A representative drop assay assessing the toxicity of Cas13 constructs. Single colonies of cells freshly transformed with the various constructs shown in (A) were grown overnight at 37°C on LB plus ampicillin and 1% glucose. The culture was then diluted 1/100 and grown until an OD_600_ of 0.4 was reached. Serial dilutions (10 μl each) of this culture were then deposited on LB plates containing 1% arabinose, which were then incubated overnight at 37°C. The graphics at bottom show the number of colony-forming units (CFU) per ml for each culture (three biological replicates) tested in plates containing 1% arabinose.

## Notes

### Competing Interest Statement

The authors have declared no competing interest.

