## Supplementary material for "Spacer acquisition in type VI CRISPR-Cas systems associated with reverse transcriptase-Cas1 fusion proteins": File S3

**Supplementary Sequences**

***Rhodovulum kholense***

*Rk* RT-Cas1

ATGCAAAATGCCGCCCCGGACCTATGGGACAGCATCACCGGCTATGATGCCCTCAACGCTGCATGGGCCCGCGTCGATGCCAATGCAGGGTCCGCCGGAGGGGATGGAGTGACGCGGCATGAGTTCCGCTCGGACCTTTTCGCCCGGCTGAACCAGCTGCGCGCCGACCTGTTGAGCGGCGAATATGTCCCGCGCCCTTACCGCAAGGTCAGCGTTCCGAAAAAGAAACCCGGCTACCGCATTCTCGCGATTCCCAGCATCCGGGACCGGGTGGTCCATACCAGCATTGCCACGGCGCTGGTGCCAATCCTCGAGCCGCATTTCGAGGAATGCTCCTTCGCATACCGACCGAACCGCGGCGTGACGAAGGCCGTCGCCCGGATCGAGCAATGGCGAAGTCGGGGCTATGAATTCGTGATCGAGGCTGACATCGTCCGCTATTTCGACAATATCGACCACGACATCCTCATGGGAAAGCTGAAGGAGCTGATTGCCGGTATCCCAGGAGCCGGGCCGGTCCTTTCGTTGACGGAACGGCTGCTGGCGCACCAGGGCAAAGGCCTCGGAACCGAAGGCGTCGGCCTGGTCCAGGGCTCGCCGCTGTCACCGCTGCTGTCCAATCTCTACCTTGATGCTCTGGACGAGGAGATCGAGGAAAGCGGCGTCAAACTGGTGCGTTTTGCCGACGACTTCGTCATCCTCTGCAAGTCGCAACGGCGGGCGGAAAAGGCCCTGGCCCACTGCGTCGAGATCCTGTCGCATCACCGTCTGCGCCTGCACGAGGAAGGCACGCGTATCGTCAATTTCGACAGAGGGTTCAACTTTATCGGCTATCTCTTCCTGAAGAACCTCGCGGTTCAAGAAAAGGCGGAACCCAAGCCAGCGGTGCCGTCAAAGCCGCTGAAATCGGAAGTGACGGACGACGGAGTGATCCTGCTCGAAGAGAAAGGGTCCAGGTTCGACCAGGGAAATCGCGTCCTTTATGTACTCGATCCCGAACACAGTCTCGGTACTCGGAATCGCAGCTTTTCGGTCCGGCGCGAGGACGGCGCCGAACTCATCGCGATCGGGCAACACCGGATTGGCCGGATCGAAGTCGGCCCGGACGTTGGCTTCGATCATGGTGCTGTCCTTCTGGCGATGGACAGCGGAACGCCGCTGGCCGCGGTCGACGGGTATGGCCAGACACGGGGCACCGTCGAACCGCGCCTCAGCCGGAAAGGCGGGCTGCACCTCGCGCAGGCGAAGGCTGTACTGACAGAGGATTTCCGGCTCTGCATCGCCCGCAGTCTTGTCGAGAGCCGGATACGGAACCAACGTACGCAGCTTTCTCGCCTCAACCGGCGACAGGGGCTGGCACCGGTGGAAGAGGCTCTTCAGGCGATGAAGCGCGAACTGGGAAAGCTAGAAACGGCCGGAAGCGTCGAAGCGGCGATGGGCCTCGAGGGGGCATCCTCGGCGCATTACTGGCGTGCGATCAGTCTCCTGGCCGGAGCCGGGGCTCCATTCAAGAGGGAACGTCCTGCCAGAAGCCCACTCAATGCCGCGATCAACTATTTAACCGGCATTCTGGAGCGTGACATACGTGCCGCGGTTCAAGGTTCCGGGCTGCATCCAGGCTTCGCATTTCTTCATGGATCACGTGACAGACATGACGGCCTGGTATTCGACATGATGGAACCGTTCCGGGCTGCAACCACAGAGGGTCTGGCGGTCTTTCTCTTCAATGCGAGGCGTCTTTCGGCCGACATGTTTTGCGAGACTGACGTCGGCAAGATAGACCTGTCGACCGAAGGGCGACAGGCGCTGGTCCAGGGATATGAGGCCGTCGTTGCCAGGCGAGTGAATCGCCCTGATCGAAAGGGCAAAATGGGGTGGAGAGCCCTCATGCTTCTCCAGTGCCGTTTTCTTGTCCGATCAATCCGCAGCGGCACATCTTCGGATTTCGTTCCATACCTCATGGAGGCATAG

*Rk* Cas2

ATGCTTCGGATCATCACTTACGACATATCCAGCGACAGGGTGAGGCGCCAAGTCGCGGCATTGCTTGAGGCGGAAGCCACGCGCGTGCAGTATTCGGTCTTCGAAGCAAGGTTATCAGATGCTGCGCTTAGACGCATCGTTTCAAGGATCGATGAGCGGCTCGCCGACAGCGATAGCCTGCGGGTGTATACGGTGGGCACTCGACATGAACGTCGCTGCCAAGTCCTGGGCTCCGGTTTGCCCGTCGATAGAGATGTGGGGTTCTGGCTGTTGTGA

*Rk* array1

AAAATTTCGGAGATCGCGCCAGTCAAGCCCGGAACCGAGGCTGAAAATGCGCCAGGACGGGATTTGCTTCAGATCGCCGCCGCCTTCCTTGTTGAAAAGCAAGGGGCTGCGGAGATCAGGGTTCTGCATGACGGTCTTTCTGACGACCGCCTGAGACTCCCTGTTGAAACGTGAGGG

*Rk* array2

ACCACAATCGGCGCCTGAAACGACGTCACGCATGCGGAATTGCCTTCAGAAAACGAGTAAGCCATTGTTCAAAAGGGAAAATATCGTGATGTGGTCAAGCATTCCGGGTTCCGCAGGCGAAAAATTCGCAGAGTCTTCTGTTTCCTTCGGAACGATGTCATATTATTCATGCCTTGTCAGCCTCCTACAGGAGCCTGGGTTCTGCATGACCGCTTTTCTGATAGCGGCCTGAGACAGCCAGCCGAACGGGTCA

*Rk* Cas13

ATGGCCAAGCATCTTCGCCCGATCAGCGAGAGCTATACCGATTTCCGTGAACCCGAAAAAGACCGCCGCCTGGGGCGCGGGCTGCACCACGACCATGAGGGGGACCAGATCCGCCCGGCGGCACTTCCCGGCTACCTTTCTGCAAACCCGAAGGCGCTCTTGCGGCAGACGATTGCCATTCTCGACAAGGTGGTCAAAAAGCGCAACCGGTTGAGCGCCATGCATCCCGAAGCCTATCGCGACAGGGTGTATCTGGCAGAGGCGCTGTGGATGCGCCTGATCTCGCGGATCAGGGAAGTCGAGAAGATCGATGAGGCAGAGATTTCCGAACTTCGCCGACTATGGAATGCCAAATCGCATCCCTATGCCATGTATGAGGTGGTTCCCTCGGGGCCGCTGAACGACAGGAACGGGCAGGCAATCCAGAAATACCGCGATGCCGATGGCCCCGAGGATCCGGTCATCGCGCGTACGGATTTCGAGGGCGTGTGGCATGGCCGGTTCTGGCAGGAGGACCCCGGGCGGGTGGATTACGGCGCCATCGCCGACCGGATCATCGCCCATCTCTTTGATCAGGAAATCGTCATCGGTGGCGGGCCCAGGCGGATAGGTTCCGTTTCGGTGCCCGAGGGCGATCCGAACGGCACGCCCGGACCGGAGACGGCACGCGGCCTGATCGTCGCGCGGGGCGAAGCCATCTCGAGAAGCGCCAGCGATCCACGAACGCCCTCAGGAAAGCTGCAGGCCCAGGCCACATGGACGATATGCGCCGAGGCCCGGTATTTCGGCGATGACGGCAAGGGCGGATCGGATATCGCAAAGGCGATCTTCGAGGAACTCGAAAAGCCGGAGAACCATGCGGGACCGATCTACCCGTCCTGGTTCGGCTCGCGCCTTTTCGACCATTTCGGCAGCTTCAAGAAAAGCCTGCCGCTGGAGAGAAAGGACCCATTCAAGGACCAGATCTGGGCGCTGCATAATGCCGTTCGGCAGTATTACCAGCGCCTCGCCAAGTCGGATCGTTTCCGCATTGCCCTCCGGAAGGCGCAGGCCGACGTGCCGGACAAGAGCGAACTTCTTTCGTTGCTTCCCAAGGATCGGCAGCATCTTCTTCTGGTCCTGGGGGGCAAGAAGAGGAATGCCGAAATGAGCGAGTTCATCCGGCTTGGCAAGCTCTTCGTCCATGCCAGCGATGCCCTTCATGACTTCAGGGAGGATGACCCTGCGGCATTGGAAGCATTCGAGAAGCGGATGACCTTTCTGGCAACCAGCATCGGCCAGAGCGAGATCAAGCAGATAGAGACCTTCGCCAGAACGTGGCGCAGCGCCGTTGGCCTGACGCTTCGCACGTTGAAAGGGCTTGCAGATCCGATGGCAAATAGGGGTAGCCCAAGAAACCGGGATATCAGCACAAAGGAAGTCATGATCGACGCTTTCAAACAGTTCGATCAGAGACATTTCGAGCGGCACATTCCGATTATCTTCGGCCTCAAGAACCTCAAAGGCGGCGACTGCCCTTCGCGCGCGTCGATATTCACGGGCAACAAGCACAAATTCACGGGCAACAAGCACAAGAGCCAGCAGGCAGAGTTGCTCTGGGCGATGATGTGCGTCGTATCTACGGTCCGCAACCGGACGAACCATTTCAATGTGCGCCGGGCCCTTATCAGATTGCTGGAAGGAGGGATCGTTCATCCGATACCGGGTGACCCCAGCAAGCTAAATGTCGCCAACCGCAAACCCCAACAGGTGTCCCCTGCTGCTCTGGATGCGTTCAAAGAGCTGCTGGAGTTCGACCTTGCAATCCAGAAGGATGCGGTGCGGACAGCCTTGGAGCAGGCCAAGGCCGACGAATTCGTCGAGATGGACAACCTCAAGCATCTGGTGGCGCATCTCGCCGGCACGGACGACATCACAGCCTTGAACCTCCCCCGGTTCGCCGCCGTCATGGACAAGGTGCGCCGCCTTGCAGATAACAATGACGTCGATCTTGCCCCTGGGCTTGTCACGCTTCAGAAGTCCCTCCCGCCGAAGGACGACCAGAAAGAGAGTGATGCGGCTCGATGCCGTTACCGGCTGCTTCTCGAACTCTACCAGTCCGGCTTTCGCGGCTGGCTGTCTGGTATTCAGTCGGACGCCGAAGGGAACAGGGAACTTTGCCATTGGGCTGTCCAACAGGTCGGGCGCTCCAAAGATGCCAGGAAAACGAAATACGATGCACGGACGAAGCGGTACTACCGCTCGCCGATCTCGCTGATCGACGATCTTGGCCTCGATCGCTACGACGACATCTCGGCGCTTCTCGATGCGTTGGCCGCGCAGTCGACGCGCGAGCAGGGGCAGGATCTGAAATACACCCCTGATGCAAGGGTTCAGAGGCAGGGGGCGAACCGGATCGAGGAGTTCCGGCAGGAACTTTTCGCACATATCTTCGCCAAGTATCTTGGGAAACCTGGCCCAGAAGACCCTGATTTCGGTTGGATCGGCGAGATCGATACCAAGCGTGACTGCTGCGACCTCGGGGAGGTCCTGAAGAGCTTCCAGCCTCCAGAACAGGCATTCCGCGCGGACTGGCACAGCCAGTTCTATGCCTGGCTCTACATGGTCCCGCCCATAGAGGCCTCACGCCTGCGCCACCAGATGCGCAAATCCGCGTCAATGGATGGCAAATGGTCCGAGACGCTTGAACTGTCCTGCGGCGAGCGCAAGGAACGCGACGAGCGGGCCCAGGGCATCCTTGCCGACATGGACCGCCTGTTCTCGCTTTACACCAAGGTTCAGGCCGCGGGCTTCAGCGGGCAGGAACACGAAGGAGCTTTGGAAGAAAAGGGACTGTTCTATCAGGACGCGTCCTTCTTCAAGGAAGTGTACCAAGAGGAAGGTACCGCCAATTATCACACGACCTTTCCCGGGACCCGCAGTGGGTTGCGTCAACTCGTCCGCTACGCCCATCTGCCGCCTCTGACGGGGATATTCGAAAAGCACAAGGTTACAGCGGACGAGGTCAAAAGCTTCGCCGGGCTTCACAACGAAAAGCGCCGCGATCCGGACAAGATCAACTGCGTAAAACAGCGGGAAGGCTTTCGGGGGGAAATCCTGCAGTCGATCAAGGACATCCCGAAGCCCAAGGACAAAAGCTATAGTAAGTTCCTTTCGGATCTGGAAGAGAAATGCCGGAATTACCAGACGGTCGCGACCGCAGCCGCCATCCATAACTTCCAGGCGAACGGGGCGCGCCTCAACGATCACGTGCGCCTCCATCAGATCGTCATGCGTGTCTTGACCCGGTTGACTGACTATGCGCTTATCTGGGAACGTGACTGCCTGTTTGCGTTCGTCGGCATGCTGTACAAGGAAACAGGCGGAACCGGGCTCAAGCCCGCCTTTGGGCCCGGGGTCAGAAAAAATGCCTTGCCGCAGATCGGCATCCTCGTGACGGCGCCGGACCTCTTGAGGAGGGCTGAAACAGAGTCAGATCACGTCTTCATCCCATATCTGGATCCCGAAACGGGCTTCGACGCGCCCAAGTTCGAAGACCGGTGTAAACTGTTGTCCGAGGTCCATTGCCAGAAGTTTCAGCTTCATTTCATAGATGCAGGTGAGCGTCCAAGTGACCAGATGCTTCGTTCGAATGAGAAGCCGGGCGGTTACGACGCATGTAAGACCTTCAAGCGGGGCACGCGCAAGCAGGACATCCGGAACGATCTGGCGCATCAGAACATTCTCGATCATCCTGGAAGGCGTGGCTTGAATCTGACCTATGTGGTCAATGCGGTCCGCTCGCTGGTGAGCTACGACCGGAAACTCAAGAATGCGGTCCCGAAATCGATCAAGCGGATCCTCGGCGAGGAAGGGCTGATCATCGAATGGCAGATGAAGGACGACCGGCTGAAGAAGCCGACGATTTATCCCGATGTGGAAACGCACCTGACAATGGTCCCGCCGCATATGATGTCCGCGCCCGTCAGCTTCACCCTGCCGAGGGCATCGGTGCGACTGGTCTCGATGACGCGGGCCATGTTCGACTTCGGCGGGTCCGGCTACGGGGACGAGGTTGTCCTGGCGGGGAAGACGGTCAAACAAGCGGTCTATCCGGATGCGTTCACGGCCCGTTTTCCGCAGACGCCGGAAACCCTGAAAAAGGAGACGAGGCTTGTTTTTACCTCAGAAAAGGAGCAAATCTCTGATTGA

*Rk* RT-Cas1_FAAA

ATGCAAAATGCCGCCCCGGACCTATGGGACAGCATCACCGGCTATGATGCCCTCAACGCTGCATGGGCCCGCGTCGATGCCAATGCAGGGTCCGCCGGAGGGGATGGAGTGACGCGGCATGAGTTCCGCTCGGACCTTTTCGCCCGGCTGAACCAGCTGCGCGCCGACCTGTTGAGCGGCGAATATGTCCCGCGCCCTTACCGCAAGGTCAGCGTTCCGAAAAAGAAACCCGGCTACCGCATTCTCGCGATTCCCAGCATCCGGGACCGGGTGGTCCATACCAGCATTGCCACGGCGCTGGTGCCAATCCTCGAGCCGCATTTCGAGGAATGCTCCTTCGCATACCGACCGAACCGCGGCGTGACGAAGGCCGTCGCCCGGATCGAGCAATGGCGAAGTCGGGGCTATGAATTCGTGATCGAGGCTGACATCGTCCGCTATTTCGACAATATCGACCACGACATCCTCATGGGAAAGCTGAAGGAGCTGATTGCCGGTATCCCAGGAGCCGGGCCGGTCCTTTCGTTGACGGAACGGCTGCTGGCGCACCAGGGCAAAGGCCTCGGAACCGAAGGCGTCGGCCTGGTCCAGGGCTCGCCGCTGTCACCGCTGCTGTCCAATCTCTACCTTGATGCTCTGGACGAGGAGATCGAGGAAAGCGGCGTCAAACTGGTGCGTTTTGCCGCGGCGTTCGTCATCCTCTGCAAGTCGCAACGGCGGGCGGAAAAGGCCCTGGCCCACTGCGTCGAGATCCTGTCGCATCACCGTCTGCGCCTGCACGAGGAAGGCACGCGTATCGTCAATTTCGACAGAGGGTTCAACTTTATCGGCTATCTCTTCCTGAAGAACCTCGCGGTTCAAGAAAAGGCGGAACCCAAGCCAGCGGTGCCGTCAAAGCCGCTGAAATCGGAAGTGACGGACGACGGAGTGATCCTGCTCGAAGAGAAAGGGTCCAGGTTCGACCAGGGAAATCGCGTCCTTTATGTACTCGATCCCGAACACAGTCTCGGTACTCGGAATCGCAGCTTTTCGGTCCGGCGCGAGGACGGCGCCGAACTCATCGCGATCGGGCAACACCGGATTGGCCGGATCGAAGTCGGCCCGGACGTTGGCTTCGATCATGGTGCTGTCCTTCTGGCGATGGACAGCGGAACGCCGCTGGCCGCGGTCGACGGGTATGGCCAGACACGGGGCACCGTCGAACCGCGCCTCAGCCGGAAAGGCGGGCTGCACCTCGCGCAGGCGAAGGCTGTACTGACAGAGGATTTCCGGCTCTGCATCGCCCGCAGTCTTGTCGAGAGCCGGATACGGAACCAACGTACGCAGCTTTCTCGCCTCAACCGGCGACAGGGGCTGGCACCGGTGGAAGAGGCTCTTCAGGCGATGAAGCGCGAACTGGGAAAGCTAGAAACGGCCGGAAGCGTCGAAGCGGCGATGGGCCTCGAGGGGGCATCCTCGGCGCATTACTGGCGTGCGATCAGTCTCCTGGCCGGAGCCGGGGCTCCATTCAAGAGGGAACGTCCTGCCAGAAGCCCACTCAATGCCGCGATCAACTATTTAACCGGCATTCTGGAGCGTGACATACGTGCCGCGGTTCAAGGTTCCGGGCTGCATCCAGGCTTCGCATTTCTTCATGGATCACGTGACAGACATGACGGCCTGGTATTCGACATGATGGAACCGTTCCGGGCTGCAACCACAGAGGGTCTGGCGGTCTTTCTCTTCAATGCGAGGCGTCTTTCGGCCGACATGTTTTGCGAGACTGACGTCGGCAAGATAGACCTGTCGACCGAAGGGCGACAGGCGCTGGTCCAGGGATATGAGGCCGTCGTTGCCAGGCGAGTGAATCGCCCTGATCGAAAGGGCAAAATGGGGTGGAGAGCCCTCATGCTTCTCCAGTGCCGTTTTCTTGTCCGATCAATCCGCAGCGGCACATCTTCGGATTTCGTTCCATACCTCATGGAGGCATAG

***Eubacterium rectale***

*Er* RT-Cas1

ATGTACTTCAAAGACGAAGACTTCGAAAAATCTATCAAAATCCTGCACCAGAAAAAAAACTCTTGCGGTATCGACAACGTTTTCATCAACCAGTTCGACGAGTTCTGGCAGTTCAACAAAGACACCATCATCTCTCAGATCAACTCTAACGCGTACAAACCGTCTGCGGTTATGCTGGAAGAAATCGTTACCAAAACCGGTAAAAAACGTCTGATCTCTCGTTACACCTGCACCGACCGTGTTATCCTGGACATCCTGAAACGTCGTCTGGTTCCGATCTTCGACAAAACCTTCTCTGACTACTCTTACGCGTACCGTGAAAACAAAGGTGTTTACGAAGCGGTTAAAAACGCGGCGAAACTGATCGAATCTGGTAAAAAATACGTTGCGGAAATCGACATCAAAGACTTCTTCGAAAACATCAACCTGCAGCGTCTGGAACACTACCTGGCGCTGAAAATCTCTGACAAAGACATGAACCAGCTGCTGCACCGTTACCTGTACATCTTCACCGTTGTTGACGACAAAAAAACCCGTAAAACCCAGGGTATCATCCAGGGTTCTTCTCTGTCTCCGCTGTTCTCTAACGTTTACATGGCGGACTTCGACAAATACCTGGAATCTAAATACTCTTTCTGCCGTTTCTCTGACAACATCAACATCTACTGCGCGTCTGAAGAAGAAGCGTACAAAGCGTTCAACGACGTTACCCACTGGCTGCAGGGTAAACTGGGTCTGAAATACAACCACGACAAATCTGGTGTTTACCCGTCTCTGGACCGTCGTTACCTGGGTTACGACTTCAAATGGCAGCGTGGTACCAACACCGTTTCTGTTTGCCGTCACAACTACGAAAAACCGAACTACTTCGGTTCTTGGCACTGCTCTGCGATCCAGAAAATCGACCGTAACTACCACATCATCAACGACGGTGTTCTGAACAAAAAAGACTTCACCATCCTGATCGAAAACGACGAACACAAAATGTACATCCCGATCGAAACCTGCACCTCTATCAACATCTACTCTAACGTTATCCTGGGTTCTTCTTTCCTGCAGTTCATCAACTCTCGTGGTCTGAACGTTAACATCTTCGACAAATACGGTAACTTCATCGGTTCTTTCCACTCTGAACACCACTACAAACGTTCTGTTACCCTGCTGAAACAGGCGTCTATCTACAACGACGAACGTGTTCGTCTGATGATCTGCGTTAAAATCGAAACCGCGTCTCTGCACAACCAGCGTGAAAACCTGCGTTACTTCAACAAACACCGTCCGTCTGGTACCCTGAAATCTGCGATCGAATACATGTCTAACTGCATCGTTGAAATGAAACAGTCTAAATCTGTTAACCAGCTGCTGACCATCGAAGCGCGTGCGAAACAGAAATACCTGCAGACCTTCGACGAAATGATCTCTGACGACCGTTTCAAATTCGACAAACGTACCCGTCGTCCGCCGCTGAACGAAGTTAACGCGATGATCTCTTTCGGTAACACCTTCATCTACCGTCGTATCGCGAACGAAATCTACCGTACCGCGCTGGACATCCGTATCGGTTTCGTTCACGCGGCGAACTCTCGTTCTGAATCTCTGAACCTGGACATCGCGGAAATCTTCAAACCGATCATCGTTGACCGTACCATCTTCACCGTTATCCACAACCTGCAGATCAACAACCGTGACCACTTCGAAAAAGAAGACAACGGTGGTATCTACCTGAACAACGCGGGTAAACGTATCTTCATCCGTGAACTGGAATACAAACTGTCTTCTAAAATCTCTATCGACGGTCAGAAACTGACCTACGACAAACTGATGAAAGCGGAAGTTTACAAAATCGTTAAATTCGTTCAGGACAACGAAAAATACAAACCGTTCAAATACACCTAA

*Er* Cas2

ATGTTCGTTATCGTTACCTACGACGTTTCTCAGAAACGTGTTACCAAAGTTATGAAAATCTGCCGTAAATACCTGAACCACATCCAGAACTCTGTTTTCGAAGGTATGATCACCGACGGTAAACTGTCTTCTCTGAAAAAAGAACTGTCTACCTGCATCGTTTGCGCGGAAGACTCTGTTGTTATCTACGAAATCCAGAACCTGAAATACACCCGTAAAGAACTGATCGGTGTTAACCGTATCAACGACAACATCTTCTAATAA

*Er* array1

AAAGTGGGTCAACGTAGTCCGTTTATGGCTTGAAGTAGAGAAAACTTGATATTATCGTATTTTGGGTGATTATAGAATAATAGGTTGACCCACTTTTTGAACCATAATTAATGATTTTTGACTAAAAAGGTGGTAATGCAATAGAACAAGTGTTATAATCAGTGGTGTGAAAGTAGCCCGATATAGAGGGCAATAACTTTTGGAGGTCGCCTTTTGAAACCTTGAATCCTAAATTCCTAA

*Er* array2

AAAGTGGGTCAACCTAGGCTTATATTGAATAAAATAAAGAAAGACAGTATTTATAAGCCTTTTGTGAATAATAATATAAACGGTTGACCCACTTTTTAATAGTAATAGGTTGATTTATGCGAAAAAAGATGATAATCTGATATAGAAAGTATTATATATACTAACGTGAATACAGCTCGATATAGTGAGCAATAAGACTTTTGCAACATTGCAACCACCAGACCAAATCAAAT

***Eubacteriaceae bacterium***

CH RT-Cas1

ATGTACCTGGACGACATCTTCACCGAAGAAAACATCCAGAACGCGCTGCACTACGTTCTGTCTCGTAAAAACTCTTGCGGTATCGACGGTATCTTCGTTAAAGACTTCGAAGAATACTGGATTCTGAACGGTCAGAAAATCCTGAAACAGGTTATGAACGGTGTTTACATGGTTTCTCCGGTTCAGCTGCGTGAAATCATCATGCCGACCGGTAAACACCGTATCATCGCGCACTACACCTGCACCGACCGTCTGATCACCCGTATCCTGGCGGAATCTCTGCAGAAAGAAGTTGACGACTCTCTGTCTGAATACTCTTACGCGTACCGTAAACAGCGTGGTGTTATCAAAGCGGTTGAACAGGCGGCGGCGTACATGCAGGCGGGTAAAATCTGGGTTCTGGAACTGGACATCGAAAACTACTTCAACAACATCAACCTGACCCTGATGGAAGAAAAAATCCGTGAAATCATCCTGGACAAAAACCTGTTCTCTCTGATGGAACAGTACCTGCGTTGCGAAGTTATGGAAGAAGAATACACCAAAACCTACATCAAAGACAAAGGTCTGGTTCAGGGTTGCTCTCTGTCTCCGGTTCTGTCTAACATCTACCTGAACAAACTGGACCAGCAGATGGAAAAAGAAGGTCTGTCTTTCTGCCGTTTCGGTGACAACATCAACATCTACTTCTACAACAAACTGGAAGCGGCGGAATGGTACGCGAAAATCAAAGCGATCATCGAAAACGAATTTGACCTGCACCTGAACATCCGTAAATCTGGTATCTACCTGGGTGTTAACCGTATCTTCCTGGGTTACTCTTTCAAAAAACAGCGTTCTGGTGAAATCCTGACCGCGCGTAACATCAAAAAAAAAACCATCTTCTACGCGAACTGGCACACCTCTGCGCTGCAGTACACCGGTAAAGAATACCACATCATCAACGACGGTATCCTGAACAAAAAAGACTACACCCTGCTGTTCGAAAACGAAAAAGAAAAAAAATACCTGCCGGTTGAAACCGTTGACAAACTGAACATCTACTCTAACGTTATCTTCGACACCGGTTTCTTCGAAATGGTTTCTCGTTACAACATCGACGTTTCTATCTTCGACAAATACGGTAAACACTGCGGTACCTTCTGCGGTTCTAAACACGCGCGTACCTCTGACATGGTTATCAAACAGGTTTCTCTGTACAACAACGACGCGAAACGTCTGTCTGTTGCGAAATCTATGCTGATCGCGGCGGCGCACAACATGCGTGCGAACGTTCGTTACTACGTTAAAAAAGAAAAACTGAAAAAAGAAAACGTTGACAAACTGTCTGCGTTCATCAAAAAACTGAACGACGCGTCTTCTATCTCTAACCTGATGATGATCGAAGCGCAGTGCCGTCAGTCTTACTACACCTACATGGGTAAAATCATCGGTGGTGGTCCGTTCCACTTCGCGCAGCGTACCAAACGTCCGCCGCGTGACGCGGTTAACGCGATGATCTCTTTCGGTAACGTTTTCCTGTACGAAAAAATCGCGACCGAAATCTACAAAACCTCTCTGGACATCAAAGTTGGTTTCCTGCACTCTACCAACCGTCGTAAAGCGACCCTGAACCTGGACATCGCGGAAATCTTCAAACCGGTTATCGTTGACCGTGTTATCTTCACCGTTATCCACAAACGTATCCTGGACGTTTCTCGTCACTTCGAATCTCAGGAAAACAACGGTGTTTACCTGAACCGTGAAGGTAAACGTCCGTTCATCAACGAACTGATGCGTAAAATGTACACCAAAATCACCGTTGACCGTAAACTGATGACCTACGAAGCGCTGATCCGTAACGAAATCTGGAAAATCTACCGTATGATCGAACGTGGTGAATCTTACAAACCGTACAAATACACCTAA

CH Cas2

ATGTTCGTTATCGTTGCATACGACGTTTCTTCTTCTCGTGGTACCAAAATCATGAAAATCTGCCGTAAATACCTGCACCACGTTCAGAAATCTGTTTTCGAAGGTACCATCACCGAAGCGAAACTGAAACAGCTGCAGCGTGAACTGAAAAACACCCTGATCCCGGACCTGGACAAAGTTTCTATCTACTGCCTGGACTCTATCAAATACGCGTCTAAAATCCAGATCGGTGCGGTTGAAGACATGGACATCTTCCTGTAGTAG

CH array1

AAAAGTCGGTCTGTCAAAAACCGCTGTTTTAAGCGCTCTAAGACGATTTTATTCTGAAAAATGGAAAAAAAGCGCCTGAAAACGGTAAAAAAATCGGTTCGTCTCAATAAGAAAAAAGTCAAATTGAGAAAACAGCTTAAATAAAGGCTCGGAGGCAGTTAAGAAAAAAGAGACAGACCGACTTTTTTCTTAGCATCAGAAAATAATATTTGACAAAGAAAAAGATGTGTGAGATAATTGTAGAAAATTAAGGAAGACAATACAAGATGTTGTTGTAGATAGCCCGATATAGAGGGCAATAAACGAGCATTATCACGGTGTTACCCGGA

CH array2

AAAAGTCGGTCTGTCAAAAACCGCTGTTTTAAGCGCTCTAAGACGATTTTGTTCTGCAAAATGAGCAAAAAACCACATGAAAACAATAAAAAAATCGGTCTGTCCCAATAAGAAAAAAGTCAAATTGAAAAAATGGTTTAAATAAAGGCTCTGAGGCAGTTGAGAAGAAAGAGACAGACCGACTTTTTTCTTAGTATCAGAAAATAATATTTGACAAAGAAAAAGATGTGTGAGATAATTGTAGGAAATTAAGGAAGACAATACAAGATATTGTTGTAGATAGCCCGATATAGAGGGCAATAAACCTTTTAACAAACTTTTCTGCATCCTA

CH Cas13

ATGAAGATAAGTAAAGAATCACACAAAAGGACGGCGGTGGCAGTGATGGAAGACCGCGTGGGTGGTGTCGTGTATGTTCCGGGTGGCTCCGGTATCGATTTGAGTAACAACCTGAAGAAAAGATCAATGGATACCAAATCACTCTACAACGTTTTCAACCAGATCCAGGCGGGCACCGCGCCGAGCGAATATGAGTGGAAGGACTACCTCTCTGAGGCGGAAAATAAAAAGCGCGAGGCCCAGAAGATGATTCAAAAAGCTAACTATGAACTGCGTCGCGAGTGCGAAGACTATGCGAAGAAAGCCAACCTGGCCGTGAGCCGTATCATCTTCTCTAAAAAACCCAAAAAAATCTTTTCCGATGATGACATTATCAGCCATATGAAGAAACAACGTCTGTCTAAGTTCAAAGGCCGTATGGAAGACTTCGTGCTAATTGCACTGCGCAAATCCCTGGTTGTGAGCACTTATAACCAAGAAGTATTTGATTCGCGTAAGGCGGCAACCGTTTTCTTGAAGAACATCGGTAAGAAGAATATTTCAGCGGACGATGAGCGTCAAATTAAACAACTGATGGCCCTTATCCGCGAGGATTATGATAAGTGGAACCCGGATAAAGATAGCTCCGACAAAAAAGAATCCTCCGGCACCAAAGTGATCCGTAGTATCGAACATCAGAACATGGTTATTCAACCGGAAAAAAATAAGTTATCTCTGTCTAAGATTAGCAATGTTGGCAAAAAGACCAAAACCAAACAGAAAGAGAAAGCTGGCCTGGACGCTTTCCTCAAGGAGTATGCGCAAATCGATGAAAACAGCCGTATGGAGTACCTGAAAAAGCTACGCCGCCTGCTGGACACCTATTTTGCAGCTCCGAGCAGCTATATAAAGGGGGCAGCGGTGTCTCTGCCGGAGAACATCAACTTTTCGTCTGAACTGAATGTTTGGGAGCGTCACGAAGCAGCGAAGAAGGTGAACATTAATTTCGTGGAAATCCCGGAAAGCTTGCTGAACGCCGAACAGAACAATAACAAGATCAACAAGGTTGAACAGGAGCACAGCCTAGAACAGCTGCGCACCGATATCAGACGTCGAAATATAACGTGCTACCACTTCGCGAATGCATTGGCAGCTGACGAGCGTTATCACACCTTGTTCTTTGAGAACATGGCAATGAATCAATTTTGGATTCATCACATGGAAAACGCGGTCGAACGTATCTTAAAAAAGTGCAACGTCGGTACTCTGTTCAAGCTGCGTATCGGCTATCTAAGCGAGAAGGTTTGGAAGGACATGCTGAACCTACTGAGCATTAAGTATATCGCGCTGGGTAAAGCGGTTTATCATTTCGCACTCGACGATATCTGGAAGGCCGATATCTGGAAGGACGCTTCCGATAAGAACAGCGGCAAGATTAACGACCTGACCCTGAAGGGCATTTCCAGCTTTGATTACGAAATGGTAAAAGCCCAAGAGGACTTGCAGCGTGAAATGGCAGTGGGTGTTGCATTCTCCACCAATAACCTGGCTCGTGTTACCTGCAAAATGGACGATCTCAGCGACGCGGAAAGCGACTTTTTGCTGTGGAATAAAGAAGCTATCCGTCGTCACGTGAAATACACCGAAAAAGGTGAGATTCTGTCAGCCATCTTACAATTTTTTGGTGGCCGCAGCCTGTGGGATGAAAGTCTGTTTGAGAAGGCATATAGCGACTCCAATTATGAACTGAAATTTCTGGACGATCTGAAGCGCGCGATTTATGCCGCTCGTAACGAAACGTTTCACTTCAAAACCGCTGCGATCGATGGCGGTTCCTGGAATACCCGTCTGTTCGGCTCGTTGTTTGAAAAAGAAGCGGGTCTCTGTCTGAATGTTGAGAAGAACAAGTTCTACTCCAACAACCTGGTGCTGTTCTACAAACAAGAGGACCTGCGTGTCTTCTTGGACAAGTTGTACGGTAAAGAGTGCAGCCGTGCGGCACAAATCCCGTCTTACAACACCATTCTACCACGTAAATCATTTAGCGACTTCATGAAGCAACTGTTAGGCCTGAAGGAACCGGTCTACGGTAGCGCCATTCTGGACCAGTGGTACTCGGCGTGTTATTACTTGTTCAAGGAGGTGTACTACAACCTGTTCCTACAGGACAGCTCCGCGAAAGCTCTTTTTGAGAAGGCTGTGAAGGCGCTGAAGGGTGCGGACAAAAAACAAGAGAAAGCAGTAGAATCTTTTCGTAAGCGCTATTGGGAGATTAGCAAGAACGCGTCTTTGGCTGAGATTTGCCAGTCTTATATCACCGAATACAATCAGCAGAACAACAAGGAGCGCAAGGTGCGCAGCGCGAACGACGGCATGTTTAACGAGCCGATTTACCAGCATTATAAGATGCTGCTGAAAGAGGCGCTGAAAATGGCGTTTGCGAGCTACATCAAAAACGATAAAGAACTGAAATTCGTCTACAAACCGACCGAAAAGTTGTTCGAGGTCTCCCAGGATAACTTCTTGCCGAATTGGAATAGCGAGAAGTACAATACCCTGATCTCTGAGGTTAAAAATAGCCCGGATCTACAGAAATGGTATATTGTTGGTAAATTCATGAATGCGCGTATGCTGAACCTGCTGTTGGGATCTATGCGCTCGTACCTGCAATATGTGAGCGACATTCAGAAGCGCGCAGCGGGTCTGGGTGAAAACCAACTGCATCTGTCCGCCGAGAACGTGGGCCAGGTGAAGAAATGGATACAAGTTCTTGAAGTCTGTCTGCTCTTGAGCGTTCGTATCTCGGATAAATTCACTGACTACTTCAAGGACGAGGAGGAATATGCGAGCTATCTGAAGGAGTATGTTGACTTCGAGGACTCGGCTATGCCGTCCGATTACTCGGCGTTGCTGGCGTTCAGCAATGAGGGTAAAATCGATTTATACGTTGATGCGTCGAACCCGAAAGTGAACCGTAATATCATTCAAGCAAAGCTGTACGCACCAGATATGGTCCTGAAGAAAGTCGTTAAGAAAATTTCACAGGACGAATGCAAAGAGTTTAACGAAAAGAAGGAACAAATCATGCAGTTTAAGAACAAAGGTGATGAAGTGAGCTGGGAAGAGCAACAGAAAATCTTGGAGTACCAAAAACTGAAGAATCGTGTTGAATTGCGTGACCTTAGCGAGTATGGCGAGCTGATCAACGAGTTGCTAGGCCAGCTGATTAACTGGAGCTACCTTCGAGAACGTGATTTACTGTACTTCCAGTTAGGTTTCCATTATTCTTGTTTGATGAACGAGTCCAAAAAGCCGGATGCGTATAAGACCATTCGCAGGGGCACGGTTAGCATCGAAAACGCAGTGCTCTACCAGATTATCGCTATGTATATTAATGGTTTTCCGGTTTACGCTCCGGAAAAGGGCGAACTGAAGCCACAGTGTAAAACGGGTAGCGCCGGTCAAAAGATTCGTGCCTTCTGCCAGTGGGCATCCATGGTTGAAAAGAAAAAATACGAGCTGTACAACGCGGGCCTGGAGCTCTTCGAAGTGGTGAAGGAGCACGATAATATTATTGACCTGCGTAATAAGATCGATCATTTTAAGTACTACCAGGGTAATGACTCTATTCTCGCGTTGTATGGTGAAATCTTCGACCGTTTTTTTACCTACGACATGAAATATCGTAATAATGTACTTAACCACCTGCAGAACATCTTGCTGCGTCACAACGTCATAATCAAACCGATTATTAGCAAGGACAAAAAAGAAGTGGGTAGAGGCAAAATGAAAGACCGTGCGGCATTTTTGCTTGAGGAGGTGAGCAGCGATCGTTTCACTTATAAGGTGAAAGAAGGTGAACGTAAAATCGACGCCAAAAACCGCCTGTACCTGGAAACCGTTCGTGATATCCTGTACTTCCCGAATCGTGCGGTCAACGATAAGGGCGAGGACGTTATCATTTGCTCCAAGAAAGCGCAGGACTTGAATGAAAAAAAGGCCGACCGCGACAAAAACCACGATAAGAGCAAGGATACGAACCAGAAAAAAGAGGGCAAGAACCAAGAGGAAAAGAGCGAGAACAAGGAGCCTTACAGCGATCGTATGACCTGGAAACCGTTTGCGGGCATCAAGTTGGAGTAA

CH RT-Cas1_FGAA

ATGTACCTGGACGACATCTTCACCGAAGAAAACATCCAGAACGCGCTGCACTACGTTCTGTCTCGTAAAAACTCTTGCGGTATCGACGGTATCTTCGTTAAAGACTTCGAAGAATACTGGATTCTGAACGGTCAGAAAATCCTGAAACAGGTTATGAACGGTGTTTACATGGTTTCTCCGGTTCAGCTGCGTGAAATCATCATGCCGACCGGTAAACACCGTATCATCGCGCACTACACCTGCACCGACCGTCTGATCACCCGTATCCTGGCGGAATCTCTGCAGAAAGAAGTTGACGACTCTCTGTCTGAATACTCTTACGCGTACCGTAAACAGCGTGGTGTTATCAAAGCGGTTGAACAGGCGGCGGCGTACATGCAGGCGGGTAAAATCTGGGTTCTGGAACTGGACATCGAAAACTACTTCAACAACATCAACCTGACCCTGATGGAAGAAAAAATCCGTGAAATCATCCTGGACAAAAACCTGTTCTCTCTGATGGAACAGTACCTGCGTTGCGAAGTTATGGAAGAAGAATACACCAAAACCTACATCAAAGACAAAGGTCTGGTTCAGGGTTGCTCTCTGTCTCCGGTTCTGTCTAACATCTACCTGAACAAACTGGACCAGCAGATGGAAAAAGAAGGTCTGTCTTTCTGCCGTTTCGGTGCCGCCATCAACATCTACTTCTACAACAAACTGGAAGCGGCGGAATGGTACGCGAAAATCAAAGCGATCATCGAAAACGAATTTGACCTGCACCTGAACATCCGTAAATCTGGTATCTACCTGGGTGTTAACCGTATCTTCCTGGGTTACTCTTTCAAAAAACAGCGTTCTGGTGAAATCCTGACCGCGCGTAACATCAAAAAAAAAACCATCTTCTACGCGAACTGGCACACCTCTGCGCTGCAGTACACCGGTAAAGAATACCACATCATCAACGACGGTATCCTGAACAAAAAAGACTACACCCTGCTGTTCGAAAACGAAAAAGAAAAAAAATACCTGCCGGTTGAAACCGTTGACAAACTGAACATCTACTCTAACGTTATCTTCGACACCGGTTTCTTCGAAATGGTTTCTCGTTACAACATCGACGTTTCTATCTTCGACAAATACGGTAAACACTGCGGTACCTTCTGCGGTTCTAAACACGCGCGTACCTCTGACATGGTTATCAAACAGGTTTCTCTGTACAACAACGACGCGAAACGTCTGTCTGTTGCGAAATCTATGCTGATCGCGGCGGCGCACAACATGCGTGCGAACGTTCGTTACTACGTTAAAAAAGAAAAACTGAAAAAAGAAAACGTTGACAAACTGTCTGCGTTCATCAAAAAACTGAACGACGCGTCTTCTATCTCTAACCTGATGATGATCGAAGCGCAGTGCCGTCAGTCTTACTACACCTACATGGGTAAAATCATCGGTGGTGGTCCGTTCCACTTCGCGCAGCGTACCAAACGTCCGCCGCGTGACGCGGTTAACGCGATGATCTCTTTCGGTAACGTTTTCCTGTACGAAAAAATCGCGACCGAAATCTACAAAACCTCTCTGGACATCAAAGTTGGTTTCCTGCACTCTACCAACCGTCGTAAAGCGACCCTGAACCTGGACATCGCGGAAATCTTCAAACCGGTTATCGTTGACCGTGTTATCTTCACCGTTATCCACAAACGTATCCTGGACGTTTCTCGTCACTTCGAATCTCAGGAAAACAACGGTGTTTACCTGAACCGTGAAGGTAAACGTCCGTTCATCAACGAACTGATGCGTAAAATGTACACCAAAATCACCGTTGACCGTAAACTGATGACCTACGAAGCGCTGATCCGTAACGAAATCTGGAAAATCTACCGTATGATCGAACGTGGTGAATCTTACAAACCGTACAAATACACCTAA

CH RT-Cas1_E543A

ATGTACCTGGACGACATCTTCACCGAAGAAAACATCCAGAACGCGCTGCACTACGTTCTGTCTCGTAAAAACTCTTGCGGTATCGACGGTATCTTCGTTAAAGACTTCGAAGAATACTGGATTCTGAACGGTCAGAAAATCCTGAAACAGGTTATGAACGGTGTTTACATGGTTTCTCCGGTTCAGCTGCGTGAAATCATCATGCCGACCGGTAAACACCGTATCATCGCGCACTACACCTGCACCGACCGTCTGATCACCCGTATCCTGGCGGAATCTCTGCAGAAAGAAGTTGACGACTCTCTGTCTGAATACTCTTACGCGTACCGTAAACAGCGTGGTGTTATCAAAGCGGTTGAACAGGCGGCGGCGTACATGCAGGCGGGTAAAATCTGGGTTCTGGAACTGGACATCGAAAACTACTTCAACAACATCAACCTGACCCTGATGGAAGAAAAAATCCGTGAAATCATCCTGGACAAAAACCTGTTCTCTCTGATGGAACAGTACCTGCGTTGCGAAGTTATGGAAGAAGAATACACCAAAACCTACATCAAAGACAAAGGTCTGGTTCAGGGTTGCTCTCTGTCTCCGGTTCTGTCTAACATCTACCTGAACAAACTGGACCAGCAGATGGAAAAAGAAGGTCTGTCTTTCTGCCGTTTCGGTGACAACATCAACATCTACTTCTACAACAAACTGGAAGCGGCGGAATGGTACGCGAAAATCAAAGCGATCATCGAAAACGAATTTGACCTGCACCTGAACATCCGTAAATCTGGTATCTACCTGGGTGTTAACCGTATCTTCCTGGGTTACTCTTTCAAAAAACAGCGTTCTGGTGAAATCCTGACCGCGCGTAACATCAAAAAAAAAACCATCTTCTACGCGAACTGGCACACCTCTGCGCTGCAGTACACCGGTAAAGAATACCACATCATCAACGACGGTATCCTGAACAAAAAAGACTACACCCTGCTGTTCGAAAACGAAAAAGAAAAAAAATACCTGCCGGTTGAAACCGTTGACAAACTGAACATCTACTCTAACGTTATCTTCGACACCGGTTTCTTCGAAATGGTTTCTCGTTACAACATCGACGTTTCTATCTTCGACAAATACGGTAAACACTGCGGTACCTTCTGCGGTTCTAAACACGCGCGTACCTCTGACATGGTTATCAAACAGGTTTCTCTGTACAACAACGACGCGAAACGTCTGTCTGTTGCGAAATCTATGCTGATCGCGGCGGCGCACAACATGCGTGCGAACGTTCGTTACTACGTTAAAAAAGAAAAACTGAAAAAAGAAAACGTTGACAAACTGTCTGCGTTCATCAAAAAACTGAACGACGCGTCTTCTATCTCTAACCTGATGATGATCGAAGCGCAGTGCCGTCAGTCTTACTACACCTACATGGGTAAAATCATCGGTGGTGGTCCGTTCCACTTCGCGCAGCGTACCAAACGTCCGCCGCGTGACGCGGTTAACGCGATGATCTCTTTCGGTAACGTTTTCCTGTACGAAAAAATCGCGACCGAAATCTACAAAACCTCTCTGGACATCAAAGTTGGTTTCCTGCACTCTACCAACCGTCGTAAAGCGACCCTGAACCTGGACATCGCGGCGATCTTCAAACCGGTTATCGTTGACCGTGTTATCTTCACCGTTATCCACAAACGTATCCTGGACGTTTCTCGTCACTTCGAATCTCAGGAAAACAACGGTGTTTACCTGAACCGTGAAGGTAAACGTCCGTTCATCAACGAACTGATGCGTAAAATGTACACCAAAATCACCGTTGACCGTAAACTGATGACCTACGAAGCGCTGATCCGTAACGAAATCTGGAAAATCTACCGTATGATCGAACGTGGTGAATCTTACAAACCGTACAAATACACCTAA

CH array1_+1A+2C

AAAAGTCGGTCTGTCAAAAACCGCTGTTTTAAGCGCTCTAAGACGATTTTATTCTGAAAAATGGAAAAAAAGCGCCTGAAAACGGTAAAAAAATCGGTTCGTCTCAATAAGAAAAAAGTCAAATTGAGAAAACAGCTTAAATAAAGGCTCGGAGGCAGTTAAGAAAAAAGAGACAGACCGACTTTTTTCTTAGCATCAGAAAATAATATTTGACAAAGAAAAAGATGTGTGAGATAATTGTAGAAAATTAAGGAAGACAATACAAGATGTTGTTACAGATAGCCCGATATAGAGGGCAATAAACGAGCATTATCACGGTGTTACCCGGA

***Human gut metagenome***

*Hm* RT-Cas1

ATGTATCTATTAAATTTCAGGAAATCATTTCCGTGCATGAATAAACTGTCTCTGAACTGCATTTCCGAGGAAAATCATCTGAAAGCGGCGTTCCAGAGCATCCTGGCGAAGGGTGCGACCGGTGGCATCGACAACATTAGCGTTACCGAATATCAGAAATCACTGTCCAAAAACATCAAACGTCTTTCCGAGAACTTGCGTGTCGGCAAGTGGAAACCGCAACCGTACCTGGGCCTGCACATCCACAAAAAGAGCGGCGGTACACGTACCCTGGGTCTGCTGAGCGTTGAAGATAAAGTGGTGCAGACTTCGATTAAGTTCTATCTTGAGCCGGTTATCGACCACATGCTGAATAGTTCAAGCTATGCCTACCGTAGCGGCCGTGGTCATGTTAAGGCTGTGAGAAGAACTCTTCACGAAAGCTCCAAGAAGTGGAATACCGTTTACCTGCGCACGGACATCAAGGACTTCTTCGATTCTATTGACCGTGATATTCTGATGAACTTGCTGTCTCGTCTGATTAAGGACACGAAATTGTTATCCCTCATTCGTTTATGCCTGACCATGGGTCGCGTGAACCCGGATATGAGCTGGCAAGAGTCGCCAGTAGGTATTCCGCAAGGCGCGATCCTCTCCCCGCTGCTCTCTAACTTGTACCTGACTAGTTTTGATACGTTTGTTAGCGGGATTACCAATGCATATATTCGCTATGCGGATGATTGTGTTTGTTGGTTTAAAGACGAGGGCTGCGCACAGGCTGCGTTGGAGGCCATGCGTGCATTTCTGCAAAACCGCCTGAAACTTCAACTGAACGACAAAGTACGCCTGGGCCGTACCGATTCTGTGCCGATGACCTATCTGGGCCTGGACTTTTATCAGGGCAAAATCTTTATCTCTGATAGCAAAAAGCAAGAACTGAGCCAGTCCATTGGCAAAATCACCATCGAGAACGGCGCACTGTCCAGCAAATATCTGAAAACCCTGGACGGTATACGTCGCTACTATGCGAAAGTTCTTCCGAATGAATTTAGCGGTATGTTCGACGTGTGGCTGAAAGACGCAATCAAGAGCTACGTCGTGAAAGGTCATATCAAGAAAAAAGAGGCATTCGCCATCTTCGGTGATATCGACGGCTATGCGGAGAAGGACTTCATCCGCAACTGGATTAGCGAGGTTGTGAAGGATACCTCCTACGCAGACGTTGAACGGAAGGTGATCGCTTCGCGTAAGCGCGAGTACCAACGTCGTGAGAGCGAAAATAGCGAGTTGATCATTAACACCCCGGGTTGTTTTTTGGGTCTCTCTGGCCGTGGCATTACCCTGCGTAAAAACGGTCAGCCAGTTCGCATCCCTCCGAGCGCGGCGTTGAAGCACATCACGATTATGAGCCAAGGTGTGTCTATGTCTAGCAATCTGATCGGCTACTGCATGGAAAACGACATCGCGATTGATTTTTTCGATCTGGGCTCCCGCCACATCGGTAGCATCCTCGCACCGAAATACATGTTCACCTCGATGTGGAAGAGCCAGATCCTGCTTGATGAGCTGCGTCGTAACGAAATCGGTCGTCGCATTATCATGGGTAAGGTCAAGAACCAGAGCTCTCTGGCTAAGTACTTCAACAAATACCATAAGACCGTCGGCGTTCAGGATGCATTCCTGCTGTATGATGACACGGTGCAAACGTTGCTGGTGAAGATCAAGAACATTAAAAATGATCAACATTTTAAATCTACCTTAATGGGCCTGGAAGCGAGCGCTGCTAGCGCGTACTGGGAATACGTTAGAGCGCTGATTGAAGACGACAATGTTGGATTTTACAGCCGCGTGAAGCAGGGTGCCACCGACCTGGTTAATAGCCTGCTGAACTATGGTTACGCTATGCTGTACCCGCGTATCTGGCAGGCGGCGTTGCGTCACAAACTGAACCCGTATATTGGCTTCGTGCATTATGCTGACAGTCAAGCCAATTTGGTTTTTGATATGATTGAGCTGTTTCGCTGCCAGTGTGTCGATCGTATTGTTATTGCGTTGATCCAGAAAAAGGAGGAGTTGAAGTTGCTGGGTGGTAAGTTAGACGAACCGACCAAAGTGAAACTGGCCAAGCACATCGCGGAGCGTTTTAACCGCCGCGAAAAATATCGTGGCGAAAGCCGACGCTTCATGGAAATCATTGACCTACAGTTTGGTGAACTGTGCGATAGCATAGCCGAGGGTGTTGCATTCCGTCCGTACTTGGCGAAGTGGTAA

*Hm* Cas2

ATGAGAGTAAGGAAGAAATTTTATGTTATAGCGTATGATACCGCAAGCGCAAAGAGACGTCGTATGATTATTAAATTGCTGGAACCATACGGCAAGCGCATCAACTACAGCGTTTACGAGTGCATGTTGACGGAGTCGCAACTGTCTCGTCTGGTTAAAGACATCTCCAAATTCGTGGTGGCCGGTAAGGACCAGGTTGCTATGTATCGTATCTGCCTGGATTGTTACGCGCATATTACCTATATCCCGGAAAAGCGCCACGACAGCGATATCGTCGTGGCGATTTAA

*Hm* Array

TATTTGCGAAAAAATAATTCTAAGACTGATCTGTTATGAACAGCAATGCATTACGAAAGAAGATGAAATTCGTGGTTTGTATTATGCTGAATACCAGATTGTTTTAGAAGCCTACATTTGAGTAGGTATGGAAACACAACCATTTTCCTATGCGCCTCAAAGCGAGAGG

***Palleniela muris***

*Pm* RT-Cas1

ATGAACACAGTATATTCAAATTTATGTCTAATGAGCACTCTGGAGCAGGCGTGGCAAATTGTTCTGCAAAAGCAGAGCGCAGGTGGTATTGACGACGTTAGCCTGGAAGACTATCGTGAGCGCCTGGGCAAGAACCTTCTCCGTCTCCAAAAGGCACTGTCAGAAAGAACATGGAAACCGCAGCCGTATATGGGTATCAGCGTTCCGAAGAACGATTCTGAGAAGCGCGAGCTGGGTCTGCTGAGCGTTGAAGACAAAATTGTGCAACAGGCGATTAAGATCCTGATTGAGCCGGTTTTCGAGAAACGTTTTTACAACTGTAGCTACGCGTACCGCGTGGGTAAGGGTCATCAGAAGGCGATCCGTCGCGTTGTTCACGAGTGCTGCCAGAAGAAAAATCAGTGGATTCTGCGACTAGACATCGACGACTTCTTCGACACCATTGACCGTGATATTTTGTTCAAACGTCTGACCGCAGCTATCCTTGATTCCGAACTCCGCCGTATTATCGAGCTTTGCGTTACGATGGGCAAAGTCGATAAAAGTATGGAATGGACCGAGCAGTGTAAAGGCATTCCGCAGGGCGCGGTGTTATCTCCGTTGCTGGCAAATTTCTACCTGACGAGCTTTGACCAGTTTGTTTGTAGCGTTACGGATGCATACGTTCGTTACGCGGATGACTTTATTGTATGGTGTGAAACCAAGGAAGAGGCAGAGAACATGCATAGCCGCATCGCGAAGCACCTGAAAAAGCACCTGCATCTGACCCTGAATGAACCGCAGATTTGTCATACCAGCGCCGGTATTGAGTACCTGGGTATCGTGGTGACGCGTAATAAGGTCTCGATCTCCCAAGAGAAACAAAACTCCATTCTGAGTCGTTTGCGCAGCATTGAAATCAAAGACAACCGTCTCAGCAAACGCTACGTGCAGTCCGTGCAGGGTGTCCACCGTTACTACGCGCGTGTTCTGCCGGAAAGCTACAGTAAAATGTTCTGCGATACGATCAAAGGGGTGATAGAGGGCTGGATTGGCAAAAACCCGAGCTACAGCGTGAAGGATTTGGAGCGTATGTTCCACGAATTACCGTTGTTTGGTAATTCCACCATCGAAGATAACAAAGAGTTGCTGCGCATCATTCGCAAGGCGAAGGCTGATGCCAAGGACGGCCGTCAAGATGTGGCCGAAGACGACAAAATGTTGAACCGTAAGCTGGTTCGTAGCCGTAAATTGGAATACCGCCGTCGCGAGAACGACAACTCGGAGCTGGTTGTTTCCAGCCATGGATATTTTATCGGTGTGTCTAATCGTGGTATCACCCTACGGAAGAACGGTAATCCTGTGTCAGTCCCACCGAGCGCGGTGCTGAAGCACATTAGCGTGATCAGCGACGGCGTGTCGATTTCGAGCAATGCAGTCAAATTCTGCATGGCAAATAATATTGCGATCGACTTCTTTGATAACCACTCCACCCACATTGCGAGCGTCGTGTCCCCGAAATACCTGCTGACCACCAATTGGCAATTGCAAGGTAGCCTGTCAGAGGAAAGACGCCTCTTCGTCGCCAAACAAATCATCATCGGCAAGCTGAAGAACCAGCTGAATCTGATGAAATACTTCAACAAGTACCATAAAAACGTTATCTCCTTCTCTCGTGAATGGTCTGACACTGAAAGATCCGTGAAAGAGGTTATCAGGAAGATTAATGACATTGATGATCTGGCGCAATATCGCATGACCCTGATGGGCTATGAAGCACAGGGTGCGGTTATCTATTGGGAATATATTCGTGATCTTATCAACGATGACGTAACTGGTTTTGAATGCCGTCAGCATCACGGCGCGACCGATTTGGTGAACTCCATGCTGAACTATGCCTATGCTATTCTTTATCCGCGTGTTTGGCAAGCGCTGTTGGTCAACAAGCTGAACCCGTATATGGGCTTCGTGCACTATCAGGAGGGTAACGCCAACCTGGTTTTTGACATGATTGAGCTGTTTCGTTCGCAAGCGGCGGATCGTGTTGTGATCTCTATGATCCAGCGCAAGGAACCATTGTCCCTGAACAACGGTATGCTGAGCGATAGCACCAAAAGCCTGCTGGCTAAAAATGTTCTGGAGCGCTTACAACGTTACGAAAAGTATCGTGGCGAAGAACGTAAATTCAGCAACATTATCGAGATGCAGGCTAAGGAGTTGGTGGATTATATGAAAACCGGCGCGTTTTATCGTCCGTACATCGCGAAGTGGTAA

*Pm* Cas2

ATGGTAATGAGGGCTAAAAAGATATATTGTGTCGTGGCCTACGACATCCGCAAAAGCAAGCGCCGTCAGCAAGTGGTGAAGCTGTTGACCCCGTTTGGTCGTCGTGTCAACAAATCTGTTTTCGAGTGCTTGTTTTCCGACGCGCAGCTGGCTCGCCTGCGTCTCGATCTGCAGAACATTATTGTTAAGAAAGAAGATCAAGTTGCGATCTACCCGATTTGTGTTAACTGCTATGCGAGAAGCGTGTATATCCCGGCTATCCGTAATGATTTCAGCGCAGTTCACGTGTTCGACTAA

*Pm* Array

TACCTTTGCGAAAAATATATACAGATGAGGTTGCAATCACTGACAATCATACAGTTATGATTGCATGATAATTTATATCATTTTCGGCTTCGTGCTGAAAATGAGGTTGTTGTAGAAGCCTACATTTGAGTAGGTATGACAACCTGCATAAGAGAACTATAGGTTAGCT

*Pm* Cas13

ATGACAAATAACAGGAATAATAAAGGAGGGAACTCCGCATACCGTAACAACAAGTCCAACCCGGATTATGAAAAACTGATTCCGGTTCCGTTTCTCGCCAACGACAAACCTGTGTGGGCAAATTACCTGAACATGGCACGTCAGAACATTTACATTACTCTTTGCCATATCACCCGTGTGCTGGGCCTGCCTTTGACTGAAAGCAGCAATCTGGAGGCCACCCTGATGCAAATTCCGGTCATTACCTTGTTGAACAAGGACAACGGCAAAGCGGAGCAGAAGGAGAAAGCGATCCGTATGCTGGATAAACACTTTCCGTTTTTGACCCCGATGGTTGAAAAGTATATTATGCTGACCTGCAAAGACCGCCAAAGCAAGGACAAAACGCCGAAAGTTTACTACGAAGTTTTTAGCATCATCCTGCCACTGATAAACCTGTTGCGCAATAAATACACCCACTATAGACTGGAGGACAAGATGCTCGATAACAAAGAAAACATCATCGACAAATCCATCTTGAAGAACATGGCTATACTCAGCCAGCTGCTCTCATACTGCTTTGTTGGTGCAACCCGTATTACGAAGGAACGCTTCGGCGCGAAATCTAACGCGGAGGGTGGTACGTTGAACGAGCGTGATTTCTATTTCCTCAACGGCTTGACCCCTCGTGAAAACCCGTGGCGATATTACCAGGCTTACAAGAAGGACAAAGAGGGCAATACCATTAAAACGCGTAAACCGAATGGCCAAGAAATCGCCACCAAAATGACCTATGAACGCGCGGACTTTAAATACGCGATTTATAACTTGAAAGATGCGAACGGTATGCCGCTGAAGGAGCGCCGTCTGACCAACGTTGGTCTGTTAATGCTGGTGTGCCTGTTTTTGGAGAAGCGCTATGCGAGCGAGTTCGCCGATCAGACAGAGTTCTTCACCCGGCGTAACACCAAAACTTATAAGCCGGAGCAAAGCGAGGTGACCATCATGCGTGAGATCATCAGCGTATATCGTATGCGTCTGCCGAAAGAGCGTATGCAATCGACCCGTGATAATTCCGCCCTGGGACTGGACATGTTGAATGAGCTGAAGAAGTGCCCGCGTCCGCTCTTTGATACTTTGTCTCCGGCGGACCAGAATCTGTTCCGCGTGGCAGTGACCGAGAATGAAAACACCCAGAACGACGAGACCATCGAGAACGGTATGATGCTAATGCTGCGTCGCACCGATCGTTTCCCGTCTTTGGCGTTGAGATTCATCGACACCAATCAGGTCTTCAAATCGATTCGATTTCAGGTGCATCTAGGCAACTATCGCTACAAGTTCTACGAAAAGAAATGGATTGATGGTAACGACGGTAAAGACAGAGTGCGTATTCTCCAGAAGGAGCTGAATGGTTTCGGCCGTCTGGACGAAATAGAGGTGACTCGCAACAACAAATGGGGTGACCTGATTCGCAAGATCGATCAGCCGCGTGAGGACACCTTTGAAACGGCACCGTATGTGACGGACCACCATGCAACCTACCTATTCAATAATAACCGTATCGGCCTGCTGTGGAATACCCAAGATCACATTGCGCTGTCCAACGGCATCTACCTGCCGTCCCTCTTGGAAAAGAACCGCATCCTGAGCGAAGACGAAAAACTGAAGGATCTGAAAGATCAGCACGGCAACGGTATTGCGGAATGTGTTGAACCGATGTGCTGGCTGAGCGTGTATGATATCCCAGCTGCGATCTTTCTGATCCACTTGCTTCACAAAAACGGTCAGACGTTCGAGTCTGCGGGTAAAACCGTTGAAGATATAATAAAAGCTACGTACAACAACTACCAACAATTCCTGAAGGGCATTGCGGGCGGAACCATTACCTCCATGGATGATTTACAAACCCAAGGTATCGGTATTAAGCCGTGTGATATCCCCGTCAAACTGCGTGAATATCTAAATGGCAAACAAACTAATATTAAAGAGGACTTCGTTTATCTGGGTTATGAGAGATTGCTGGACATGTATGAACGTAATGAAAAACGACTGGAGCGTTTCCAACGTGATAACGACACATACGGTAGCAAAGAAAACAAAATCGGCAAGAAGGCTTACATCGACATCCGTCCGGGTACTCTGGCTCGTCATATTGCTGATGATATCATGTTTTTTGTTCCGACGGACAAGAATAACAACAAGGTCCGGATTACCGGTCAAAACTTTAATATCATGCAAGCGGCGATTGCTCAGTACACCGGCAATATCGTGCAACTGCGTACCCTACTGCAAAAAGGCGAAATTATCAATTGTGAACGCCAGCATCCGTTCATGCACAAAGTGCTGGCGACTAAGCCGTGCAGCTTCGCAGCCCTGTACACGGCCTATCTGAAGGCGAAACGCGCATATTTGGCCAACCTCATCAAGGAATACCAAAAGGCGCGTAAAAAGACCAATGATATAAAGACCCTGAAGAAGCAGTTTGTGAAGGACTACAGCAAATTAGCGGCGCTTTCATTTCTGCACGTTGACCGCCAGAAGTGGGCGGAGCGTAACGAAGATTACTATAAACAGCTTGCCGAGAGATATGCGACGATCGAGCTGCCGCATGGTATTTTTCTGAACGCTATCCATGATGCGATTATGCAGCTGCCGGAGTCTAAGAATATGGACATTGATAAAGAAAAGGACAACGTGAGCGGCCTGATTCTGAAGTACTTCAAGGCTAGCGGTGACGCTAGTCAGCCGTTCTATGGCAACAAATTTAAGCGCAACTACAAATACTTGGACATGATGGTCAAACCGGAGTGGAGCTACTCGGGTCCGCAGCCGAGCCTGGAGAAGCACTACTATTCTATCAGCGAAATGGCGAACATGCAGAAGCAGTGGGGTCAGACCAAAGAGCTGCGCGAGAAGCGTGTTGCAGTCTATGCCAAAGCGGTGGATAAAAAAACTAACCATCTGCTGGAGAAAGCGCGTGCTAAGTTTGGGAATGACATCAGAAAGCTGGAGCGTCGTGAAAAATCGATTCAGCGTGAATGTAATGAAGAAAAAAAGGCCGCGATTGAAAAGCTCCAACATTGCCGTACCTCCTACTCCGCGAATGAGCGCGAAATCCGTCGTTACAAGGTTCAGGATATCTTGATTTTCCTAATGGCGATCGACCACCTGACCGCGGCGATGGGTACCAAAGGTAATTTCAGCTCTTACAAGTTGCGGAACATCGGCTGCGCGAGCGAGAGCTCGGATATCCTGTCTATGCTGATGCCATTCTCTATGACGCTAGAGATTCCGAGCCTGATTAAGGACGAGCCGGCAAAGGTCATCACCATCAAGCAAGATGATTTAAAACTGAAAAACTACGGTGACTTCTTTCGTTTTATCTACGATCGTCGCATCCGTACCCTGCTGCAAAATGTTAAGGAAAACGAGTTCACCCGTCAGCAAATTGAAGAAGAACTGGAGTGCTATAATCGTATGCGGATTCCAGTGTTTAAGCAAGTTCTGTCCACCGAGGAGAAGATTTGGAAACAGTTGCCGAGTGAGCAGCTGCACAAAAAGGCGGACGACAAAGAACACCCGGTTAATATTGACTTCAAGTATTTAATGCAGTTCGTTAACATTAAAGAAAGCAATATCGAGATCATAAAAGCGATCCGCAACGCATTTAGCCATAACCAATACCCGACCGACAGCGCTGTAGCTCGTCTTGTGTTCGATAAGAAGCAGATCCCGGAGCTGGCCAAAGAGATTACCAAGTTGTTACACAACAAAACCAACCAGATTAATTTAAAAGACATGGAAGATAAGCAGTAA

*Pm* RT-Cas1_E665A

ATGAACACAGTATATTCAAATTTATGTCTAATGAGCACTCTGGAGCAGGCGTGGCAAATTGTTCTGCAAAAGCAGAGCGCAGGTGGTATTGACGACGTTAGCCTGGAAGACTATCGTGAGCGCCTGGGCAAGAACCTTCTCCGTCTCCAAAAGGCACTGTCAGAAAGAACATGGAAACCGCAGCCGTATATGGGTATCAGCGTTCCGAAGAACGATTCTGAGAAGCGCGAGCTGGGTCTGCTGAGCGTTGAAGACAAAATTGTGCAACAGGCGATTAAGATCCTGATTGAGCCGGTTTTCGAGAAACGTTTTTACAACTGTAGCTACGCGTACCGCGTGGGTAAGGGTCATCAGAAGGCGATCCGTCGCGTTGTTCACGAGTGCTGCCAGAAGAAAAATCAGTGGATTCTGCGACTAGACATCGACGACTTCTTCGACACCATTGACCGTGATATTTTGTTCAAACGTCTGACCGCAGCTATCCTTGATTCCGAACTCCGCCGTATTATCGAGCTTTGCGTTACGATGGGCAAAGTCGATAAAAGTATGGAATGGACCGAGCAGTGTAAAGGCATTCCGCAGGGCGCGGTGTTATCTCCGTTGCTGGCAAATTTCTACCTGACGAGCTTTGACCAGTTTGTTTGTAGCGTTACGGATGCATACGTTCGTTACGCGGATGACTTTATTGTATGGTGTGAAACCAAGGAAGAGGCAGAGAACATGCATAGCCGCATCGCGAAGCACCTGAAAAAGCACCTGCATCTGACCCTGAATGAACCGCAGATTTGTCATACCAGCGCCGGTATTGAGTACCTGGGTATCGTGGTGACGCGTAATAAGGTCTCGATCTCCCAAGAGAAACAAAACTCCATTCTGAGTCGTTTGCGCAGCATTGAAATCAAAGACAACCGTCTCAGCAAACGCTACGTGCAGTCCGTGCAGGGTGTCCACCGTTACTACGCGCGTGTTCTGCCGGAAAGCTACAGTAAAATGTTCTGCGATACGATCAAAGGGGTGATAGAGGGCTGGATTGGCAAAAACCCGAGCTACAGCGTGAAGGATTTGGAGCGTATGTTCCACGAATTACCGTTGTTTGGTAATTCCACCATCGAAGATAACAAAGAGTTGCTGCGCATCATTCGCAAGGCGAAGGCTGATGCCAAGGACGGCCGTCAAGATGTGGCCGAAGACGACAAAATGTTGAACCGTAAGCTGGTTCGTAGCCGTAAATTGGAATACCGCCGTCGCGAGAACGACAACTCGGAGCTGGTTGTTTCCAGCCATGGATATTTTATCGGTGTGTCTAATCGTGGTATCACCCTACGGAAGAACGGTAATCCTGTGTCAGTCCCACCGAGCGCGGTGCTGAAGCACATTAGCGTGATCAGCGACGGCGTGTCGATTTCGAGCAATGCAGTCAAATTCTGCATGGCAAATAATATTGCGATCGACTTCTTTGATAACCACTCCACCCACATTGCGAGCGTCGTGTCCCCGAAATACCTGCTGACCACCAATTGGCAATTGCAAGGTAGCCTGTCAGAGGAAAGACGCCTCTTCGTCGCCAAACAAATCATCATCGGCAAGCTGAAGAACCAGCTGAATCTGATGAAATACTTCAACAAGTACCATAAAAACGTTATCTCCTTCTCTCGTGAATGGTCTGACACTGAAAGATCCGTGAAAGAGGTTATCAGGAAGATTAATGACATTGATGATCTGGCGCAATATCGCATGACCCTGATGGGCTATGAAGCACAGGGTGCGGTTATCTATTGGGAATATATTCGTGATCTTATCAACGATGACGTAACTGGTTTTGAATGCCGTCAGCATCACGGCGCGACCGATTTGGTGAACTCCATGCTGAACTATGCCTATGCTATTCTTTATCCGCGTGTTTGGCAAGCGCTGTTGGTCAACAAGCTGAACCCGTATATGGGCTTCGTGCACTATCAGGAGGGTAACGCCAACCTGGTTTTTGACATGATTGCGCTGTTTCGTTCGCAAGCGGCGGATCGTGTTGTGATCTCTATGATCCAGCGCAAGGAACCATTGTCCCTGAACAACGGTATGCTGAGCGATAGCACCAAAAGCCTGCTGGCTAAAAATGTTCTGGAGCGCTTACAACGTTACGAAAAGTATCGTGGCGAAGAACGTAAATTCAGCAACATTATCGAGATGCAGGCTAAGGAGTTGGTGGATTATATGAAAACCGGCGCGTTTTATCGTCCGTACATCGCGAAGTGGTAA

*Pm* RT-Cas1_YAAA

ATGAACACAGTATATTCAAATTTATGTCTAATGAGCACTCTGGAGCAGGCGTGGCAAATTGTTCTGCAAAAGCAGAGCGCAGGTGGTATTGACGACGTTAGCCTGGAAGACTATCGTGAGCGCCTGGGCAAGAACCTTCTCCGTCTCCAAAAGGCACTGTCAGAAAGAACATGGAAACCGCAGCCGTATATGGGTATCAGCGTTCCGAAGAACGATTCTGAGAAGCGCGAGCTGGGTCTGCTGAGCGTTGAAGACAAAATTGTGCAACAGGCGATTAAGATCCTGATTGAGCCGGTTTTCGAGAAACGTTTTTACAACTGTAGCTACGCGTACCGCGTGGGTAAGGGTCATCAGAAGGCGATCCGTCGCGTTGTTCACGAGTGCTGCCAGAAGAAAAATCAGTGGATTCTGCGACTAGACATCGACGACTTCTTCGACACCATTGACCGTGATATTTTGTTCAAACGTCTGACCGCAGCTATCCTTGATTCCGAACTCCGCCGTATTATCGAGCTTTGCGTTACGATGGGCAAAGTCGATAAAAGTATGGAATGGACCGAGCAGTGTAAAGGCATTCCGCAGGGCGCGGTGTTATCTCCGTTGCTGGCAAATTTCTACCTGACGAGCTTTGACCAGTTTGTTTGTAGCGTTACGGATGCATACGTTCGTTACGCGGCTGCCTTTATTGTATGGTGTGAAACCAAGGAAGAGGCAGAGAACATGCATAGCCGCATCGCGAAGCACCTGAAAAAGCACCTGCATCTGACCCTGAATGAACCGCAGATTTGTCATACCAGCGCCGGTATTGAGTACCTGGGTATCGTGGTGACGCGTAATAAGGTCTCGATCTCCCAAGAGAAACAAAACTCCATTCTGAGTCGTTTGCGCAGCATTGAAATCAAAGACAACCGTCTCAGCAAACGCTACGTGCAGTCCGTGCAGGGTGTCCACCGTTACTACGCGCGTGTTCTGCCGGAAAGCTACAGTAAAATGTTCTGCGATACGATCAAAGGGGTGATAGAGGGCTGGATTGGCAAAAACCCGAGCTACAGCGTGAAGGATTTGGAGCGTATGTTCCACGAATTACCGTTGTTTGGTAATTCCACCATCGAAGATAACAAAGAGTTGCTGCGCATCATTCGCAAGGCGAAGGCTGATGCCAAGGACGGCCGTCAAGATGTGGCCGAAGACGACAAAATGTTGAACCGTAAGCTGGTTCGTAGCCGTAAATTGGAATACCGCCGTCGCGAGAACGACAACTCGGAGCTGGTTGTTTCCAGCCATGGATATTTTATCGGTGTGTCTAATCGTGGTATCACCCTACGGAAGAACGGTAATCCTGTGTCAGTCCCACCGAGCGCGGTGCTGAAGCACATTAGCGTGATCAGCGACGGCGTGTCGATTTCGAGCAATGCAGTCAAATTCTGCATGGCAAATAATATTGCGATCGACTTCTTTGATAACCACTCCACCCACATTGCGAGCGTCGTGTCCCCGAAATACCTGCTGACCACCAATTGGCAATTGCAAGGTAGCCTGTCAGAGGAAAGACGCCTCTTCGTCGCCAAACAAATCATCATCGGCAAGCTGAAGAACCAGCTGAATCTGATGAAATACTTCAACAAGTACCATAAAAACGTTATCTCCTTCTCTCGTGAATGGTCTGACACTGAAAGATCCGTGAAAGAGGTTATCAGGAAGATTAATGACATTGATGATCTGGCGCAATATCGCATGACCCTGATGGGCTATGAAGCACAGGGTGCGGTTATCTATTGGGAATATATTCGTGATCTTATCAACGATGACGTAACTGGTTTTGAATGCCGTCAGCATCACGGCGCGACCGATTTGGTGAACTCCATGCTGAACTATGCCTATGCTATTCTTTATCCGCGTGTTTGGCAAGCGCTGTTGGTCAACAAGCTGAACCCGTATATGGGCTTCGTGCACTATCAGGAGGGTAACGCCAACCTGGTTTTTGACATGATTGAGCTGTTTCGTTCGCAAGCGGCGGATCGTGTTGTGATCTCTATGATCCAGCGCAAGGAACCATTGTCCCTGAACAACGGTATGCTGAGCGATAGCACCAAAAGCCTGCTGGCTAAAAATGTTCTGGAGCGCTTACAACGTTACGAAAAGTATCGTGGCGAAGAACGTAAATTCAGCAACATTATCGAGATGCAGGCTAAGGAGTTGGTGGATTATATGAAAACCGGCGCGTTTTATCGTCCGTACATCGCGAAGTGGTAA

*Pm* Cas13_D1D2 (HEPN1-HEPN2)

ATGACAAATAACAGGAATAATAAAGGAGGGAACTCCGCATACCGTAACAACAAGTCCAACCCGGATTATGAAAAACTGATTCCGGTTCCGTTTCTCGCCAACGACAAACCTGTGTGGGCAAATTACCTGAACATGGCACGTCAGAACATTTACATTACTCTTTGCCATATCACCCGTGTGCTGGGCCTGCCTTTGACTGAAAGCAGCAATCTGGAGGCCACCCTGATGCAAATTCCGGTCATTACCTTGTTGAACAAGGACAACGGCAAAGCGGAGCAGAAGGAGAAAGCGATCCGTATGCTGGATAAACACTTTCCGTTTTTGACCCCGATGGTTGAAAAGTATATTATGCTGACCTGCAAAGACCGCCAAAGCAAGGACAAAACGCCGAAAGTTTACTACGAAGTTTTTAGCATCATCCTGCCACTGATAAACCTGTTGGCCAATAAATACACCGCCTATAGACTGGAGGACAAGATGCTCGATAACAAAGAAAACATCATCGACAAATCCATCTTGAAGAACATGGCTATACTCAGCCAGCTGCTCTCATACTGCTTTGTTGGTGCAACCCGTATTACGAAGGAACGCTTCGGCGCGAAATCTAACGCGGAGGGTGGTACGTTGAACGAGCGTGATTTCTATTTCCTCAACGGCTTGACCCCTCGTGAAAACCCGTGGCGATATTACCAGGCTTACAAGAAGGACAAAGAGGGCAATACCATTAAAACGCGTAAACCGAATGGCCAAGAAATCGCCACCAAAATGACCTATGAACGCGCGGACTTTAAATACGCGATTTATAACTTGAAAGATGCGAACGGTATGCCGCTGAAGGAGCGCCGTCTGACCAACGTTGGTCTGTTAATGCTGGTGTGCCTGTTTTTGGAGAAGCGCTATGCGAGCGAGTTCGCCGATCAGACAGAGTTCTTCACCCGGCGTAACACCAAAACTTATAAGCCGGAGCAAAGCGAGGTGACCATCATGCGTGAGATCATCAGCGTATATCGTATGCGTCTGCCGAAAGAGCGTATGCAATCGACCCGTGATAATTCCGCCCTGGGACTGGACATGTTGAATGAGCTGAAGAAGTGCCCGCGTCCGCTCTTTGATACTTTGTCTCCGGCGGACCAGAATCTGTTCCGCGTGGCAGTGACCGAGAATGAAAACACCCAGAACGACGAGACCATCGAGAACGGTATGATGCTAATGCTGCGTCGCACCGATCGTTTCCCGTCTTTGGCGTTGAGATTCATCGACACCAATCAGGTCTTCAAATCGATTCGATTTCAGGTGCATCTAGGCAACTATCGCTACAAGTTCTACGAAAAGAAATGGATTGATGGTAACGACGGTAAAGACAGAGTGCGTATTCTCCAGAAGGAGCTGAATGGTTTCGGCCGTCTGGACGAAATAGAGGTGACTCGCAACAACAAATGGGGTGACCTGATTCGCAAGATCGATCAGCCGCGTGAGGACACCTTTGAAACGGCACCGTATGTGACGGACCACCATGCAACCTACCTATTCAATAATAACCGTATCGGCCTGCTGTGGAATACCCAAGATCACATTGCGCTGTCCAACGGCATCTACCTGCCGTCCCTCTTGGAAAAGAACCGCATCCTGAGCGAAGACGAAAAACTGAAGGATCTGAAAGATCAGCACGGCAACGGTATTGCGGAATGTGTTGAACCGATGTGCTGGCTGAGCGTGTATGATATCCCAGCTGCGATCTTTCTGATCCACTTGCTTCACAAAAACGGTCAGACGTTCGAGTCTGCGGGTAAAACCGTTGAAGATATAATAAAAGCTACGTACAACAACTACCAACAATTCCTGAAGGGCATTGCGGGCGGAACCATTACCTCCATGGATGATTTACAAACCCAAGGTATCGGTATTAAGCCGTGTGATATCCCCGTCAAACTGCGTGAATATCTAAATGGCAAACAAACTAATATTAAAGAGGACTTCGTTTATCTGGGTTATGAGAGATTGCTGGACATGTATGAACGTAATGAAAAACGACTGGAGCGTTTCCAACGTGATAACGACACATACGGTAGCAAAGAAAACAAAATCGGCAAGAAGGCTTACATCGACATCCGTCCGGGTACTCTGGCTCGTCATATTGCTGATGATATCATGTTTTTTGTTCCGACGGACAAGAATAACAACAAGGTCCGGATTACCGGTCAAAACTTTAATATCATGCAAGCGGCGATTGCTCAGTACACCGGCAATATCGTGCAACTGCGTACCCTACTGCAAAAAGGCGAAATTATCAATTGTGAACGCCAGCATCCGTTCATGCACAAAGTGCTGGCGACTAAGCCGTGCAGCTTCGCAGCCCTGTACACGGCCTATCTGAAGGCGAAACGCGCATATTTGGCCAACCTCATCAAGGAATACCAAAAGGCGCGTAAAAAGACCAATGATATAAAGACCCTGAAGAAGCAGTTTGTGAAGGACTACAGCAAATTAGCGGCGCTTTCATTTCTGCACGTTGACCGCCAGAAGTGGGCGGAGCGTAACGAAGATTACTATAAACAGCTTGCCGAGAGATATGCGACGATCGAGCTGCCGCATGGTATTTTTCTGAACGCTATCCATGATGCGATTATGCAGCTGCCGGAGTCTAAGAATATGGACATTGATAAAGAAAAGGACAACGTGAGCGGCCTGATTCTGAAGTACTTCAAGGCTAGCGGTGACGCTAGTCAGCCGTTCTATGGCAACAAATTTAAGCGCAACTACAAATACTTGGACATGATGGTCAAACCGGAGTGGAGCTACTCGGGTCCGCAGCCGAGCCTGGAGAAGCACTACTATTCTATCAGCGAAATGGCGAACATGCAGAAGCAGTGGGGTCAGACCAAAGAGCTGCGCGAGAAGCGTGTTGCAGTCTATGCCAAAGCGGTGGATAAAAAAACTAACCATCTGCTGGAGAAAGCGCGTGCTAAGTTTGGGAATGACATCAGAAAGCTGGAGCGTCGTGAAAAATCGATTCAGCGTGAATGTAATGAAGAAAAAAAGGCCGCGATTGAAAAGCTCCAACATTGCCGTACCTCCTACTCCGCGAATGAGCGCGAAATCCGTCGTTACAAGGTTCAGGATATCTTGATTTTCCTAATGGCGATCGACCACCTGACCGCGGCGATGGGTACCAAAGGTAATTTCAGCTCTTACAAGTTGCGGAACATCGGCTGCGCGAGCGAGAGCTCGGATATCCTGTCTATGCTGATGCCATTCTCTATGACGCTAGAGATTCCGAGCCTGATTAAGGACGAGCCGGCAAAGGTCATCACCATCAAGCAAGATGATTTAAAACTGAAAAACTACGGTGACTTCTTTCGTTTTATCTACGATCGTCGCATCCGTACCCTGCTGCAAAATGTTAAGGAAAACGAGTTCACCCGTCAGCAAATTGAAGAAGAACTGGAGTGCTATAATCGTATGCGGATTCCAGTGTTTAAGCAAGTTCTGTCCACCGAGGAGAAGATTTGGAAACAGTTGCCGAGTGAGCAGCTGCACAAAAAGGCGGACGACAAAGAACACCCGGTTAATATTGACTTCAAGTATTTAATGCAGTTCGTTAACATTAAAGAAAGCAATATCGAGATCATAAAAGCGATCGCCAACGCATTTAGCGCTAACCAATACCCGACCGACAGCGCTGTAGCTCGTCTTGTGTTCGATAAGAAGCAGATCCCGGAGCTGGCCAAAGAGATTACCAAGTTGTTACACAACAAAACCAACCAGATTAATTTAAAAGACATGGAAGATAAGCAGTAA

*Pm* Cas13_KK (K444A-K445A)

ATGACAAATAACAGGAATAATAAAGGAGGGAACTCCGCATACCGTAACAACAAGTCCAACCCGGATTATGAAAAACTGATTCCGGTTCCGTTTCTCGCCAACGACAAACCTGTGTGGGCAAATTACCTGAACATGGCACGTCAGAACATTTACATTACTCTTTGCCATATCACCCGTGTGCTGGGCCTGCCTTTGACTGAAAGCAGCAATCTGGAGGCCACCCTGATGCAAATTCCGGTCATTACCTTGTTGAACAAGGACAACGGCAAAGCGGAGCAGAAGGAGAAAGCGATCCGTATGCTGGATAAACACTTTCCGTTTTTGACCCCGATGGTTGAAAAGTATATTATGCTGACCTGCAAAGACCGCCAAAGCAAGGACAAAACGCCGAAAGTTTACTACGAAGTTTTTAGCATCATCCTGCCACTGATAAACCTGTTGCGCAATAAATACACCCACTATAGACTGGAGGACAAGATGCTCGATAACAAAGAAAACATCATCGACAAATCCATCTTGAAGAACATGGCTATACTCAGCCAGCTGCTCTCATACTGCTTTGTTGGTGCAACCCGTATTACGAAGGAACGCTTCGGCGCGAAATCTAACGCGGAGGGTGGTACGTTGAACGAGCGTGATTTCTATTTCCTCAACGGCTTGACCCCTCGTGAAAACCCGTGGCGATATTACCAGGCTTACAAGAAGGACAAAGAGGGCAATACCATTAAAACGCGTAAACCGAATGGCCAAGAAATCGCCACCAAAATGACCTATGAACGCGCGGACTTTAAATACGCGATTTATAACTTGAAAGATGCGAACGGTATGCCGCTGAAGGAGCGCCGTCTGACCAACGTTGGTCTGTTAATGCTGGTGTGCCTGTTTTTGGAGAAGCGCTATGCGAGCGAGTTCGCCGATCAGACAGAGTTCTTCACCCGGCGTAACACCAAAACTTATAAGCCGGAGCAAAGCGAGGTGACCATCATGCGTGAGATCATCAGCGTATATCGTATGCGTCTGCCGAAAGAGCGTATGCAATCGACCCGTGATAATTCCGCCCTGGGACTGGACATGTTGAATGAGCTGAAGAAGTGCCCGCGTCCGCTCTTTGATACTTTGTCTCCGGCGGACCAGAATCTGTTCCGCGTGGCAGTGACCGAGAATGAAAACACCCAGAACGACGAGACCATCGAGAACGGTATGATGCTAATGCTGCGTCGCACCGATCGTTTCCCGTCTTTGGCGTTGAGATTCATCGACACCAATCAGGTCTTCAAATCGATTCGATTTCAGGTGCATCTAGGCAACTATCGCTACAAGTTCTACGAAGCGGCATGGATTGATGGTAACGACGGTAAAGACAGAGTGCGTATTCTCCAGAAGGAGCTGAATGGTTTCGGCCGTCTGGACGAAATAGAGGTGACTCGCAACAACAAATGGGGTGACCTGATTCGCAAGATCGATCAGCCGCGTGAGGACACCTTTGAAACGGCACCGTATGTGACGGACCACCATGCAACCTACCTATTCAATAATAACCGTATCGGCCTGCTGTGGAATACCCAAGATCACATTGCGCTGTCCAACGGCATCTACCTGCCGTCCCTCTTGGAAAAGAACCGCATCCTGAGCGAAGACGAAAAACTGAAGGATCTGAAAGATCAGCACGGCAACGGTATTGCGGAATGTGTTGAACCGATGTGCTGGCTGAGCGTGTATGATATCCCAGCTGCGATCTTTCTGATCCACTTGCTTCACAAAAACGGTCAGACGTTCGAGTCTGCGGGTAAAACCGTTGAAGATATAATAAAAGCTACGTACAACAACTACCAACAATTCCTGAAGGGCATTGCGGGCGGAACCATTACCTCCATGGATGATTTACAAACCCAAGGTATCGGTATTAAGCCGTGTGATATCCCCGTCAAACTGCGTGAATATCTAAATGGCAAACAAACTAATATTAAAGAGGACTTCGTTTATCTGGGTTATGAGAGATTGCTGGACATGTATGAACGTAATGAAAAACGACTGGAGCGTTTCCAACGTGATAACGACACATACGGTAGCAAAGAAAACAAAATCGGCAAGAAGGCTTACATCGACATCCGTCCGGGTACTCTGGCTCGTCATATTGCTGATGATATCATGTTTTTTGTTCCGACGGACAAGAATAACAACAAGGTCCGGATTACCGGTCAAAACTTTAATATCATGCAAGCGGCGATTGCTCAGTACACCGGCAATATCGTGCAACTGCGTACCCTACTGCAAAAAGGCGAAATTATCAATTGTGAACGCCAGCATCCGTTCATGCACAAAGTGCTGGCGACTAAGCCGTGCAGCTTCGCAGCCCTGTACACGGCCTATCTGAAGGCGAAACGCGCATATTTGGCCAACCTCATCAAGGAATACCAAAAGGCGCGTAAAAAGACCAATGATATAAAGACCCTGAAGAAGCAGTTTGTGAAGGACTACAGCAAATTAGCGGCGCTTTCATTTCTGCACGTTGACCGCCAGAAGTGGGCGGAGCGTAACGAAGATTACTATAAACAGCTTGCCGAGAGATATGCGACGATCGAGCTGCCGCATGGTATTTTTCTGAACGCTATCCATGATGCGATTATGCAGCTGCCGGAGTCTAAGAATATGGACATTGATAAAGAAAAGGACAACGTGAGCGGCCTGATTCTGAAGTACTTCAAGGCTAGCGGTGACGCTAGTCAGCCGTTCTATGGCAACAAATTTAAGCGCAACTACAAATACTTGGACATGATGGTCAAACCGGAGTGGAGCTACTCGGGTCCGCAGCCGAGCCTGGAGAAGCACTACTATTCTATCAGCGAAATGGCGAACATGCAGAAGCAGTGGGGTCAGACCAAAGAGCTGCGCGAGAAGCGTGTTGCAGTCTATGCCAAAGCGGTGGATAAAAAAACTAACCATCTGCTGGAGAAAGCGCGTGCTAAGTTTGGGAATGACATCAGAAAGCTGGAGCGTCGTGAAAAATCGATTCAGCGTGAATGTAATGAAGAAAAAAAGGCCGCGATTGAAAAGCTCCAACATTGCCGTACCTCCTACTCCGCGAATGAGCGCGAAATCCGTCGTTACAAGGTTCAGGATATCTTGATTTTCCTAATGGCGATCGACCACCTGACCGCGGCGATGGGTACCAAAGGTAATTTCAGCTCTTACAAGTTGCGGAACATCGGCTGCGCGAGCGAGAGCTCGGATATCCTGTCTATGCTGATGCCATTCTCTATGACGCTAGAGATTCCGAGCCTGATTAAGGACGAGCCGGCAAAGGTCATCACCATCAAGCAAGATGATTTAAAACTGAAAAACTACGGTGACTTCTTTCGTTTTATCTACGATCGTCGCATCCGTACCCTGCTGCAAAATGTTAAGGAAAACGAGTTCACCCGTCAGCAAATTGAAGAAGAACTGGAGTGCTATAATCGTATGCGGATTCCAGTGTTTAAGCAAGTTCTGTCCACCGAGGAGAAGATTTGGAAACAGTTGCCGAGTGAGCAGCTGCACAAAAAGGCGGACGACAAAGAACACCCGGTTAATATTGACTTCAAGTATTTAATGCAGTTCGTTAACATTAAAGAAAGCAATATCGAGATCATAAAAGCGATCCGCAACGCATTTAGCCATAACCAATACCCGACCGACAGCGCTGTAGCTCGTCTTGTGTTCGATAAGAAGCAGATCCCGGAGCTGGCCAAAGAGATTACCAAGTTGTTACACAACAAAACCAACCAGATTAATTTAAAAGACATGGAAGATAAGCAGTAA

*Pm* Cas13_Δ210C

ATGACAAATAACAGGAATAATAAAGGAGGGAACTCCGCATACCGTAACAACAAGTCCAACCCGGATTATGAAAAACTGATTCCGGTTCCGTTTCTCGCCAACGACAAACCTGTGTGGGCAAATTACCTGAACATGGCACGTCAGAACATTTACATTACTCTTTGCCATATCACCCGTGTGCTGGGCCTGCCTTTGACTGAAAGCAGCAATCTGGAGGCCACCCTGATGCAAATTCCGGTCATTACCTTGTTGAACAAGGACAACGGCAAAGCGGAGCAGAAGGAGAAAGCGATCCGTATGCTGGATAAACACTTTCCGTTTTTGACCCCGATGGTTGAAAAGTATATTATGCTGACCTGCAAAGACCGCCAAAGCAAGGACAAAACGCCGAAAGTTTACTACGAAGTTTTTAGCATCATCCTGCCACTGATAAACCTGTTGCGCAATAAATACACCCACTATAGACTGGAGGACAAGATGCTCGATAACAAAGAAAACATCATCGACAAATCCATCTTGAAGAACATGGCTATACTCAGCCAGCTGCTCTCATACTGCTTTGTTGGTGCAACCCGTATTACGAAGGAACGCTTCGGCGCGAAATCTAACGCGGAGGGTGGTACGTTGAACGAGCGTGATTTCTATTTCCTCAACGGCTTGACCCCTCGTGAAAACCCGTGGCGATATTACCAGGCTTACAAGAAGGACAAAGAGGGCAATACCATTAAAACGCGTAAACCGAATGGCCAAGAAATCGCCACCAAAATGACCTATGAACGCGCGGACTTTAAATACGCGATTTATAACTTGAAAGATGCGAACGGTATGCCGCTGAAGGAGCGCCGTCTGACCAACGTTGGTCTGTTAATGCTGGTGTGCCTGTTTTTGGAGAAGCGCTATGCGAGCGAGTTCGCCGATCAGACAGAGTTCTTCACCCGGCGTAACACCAAAACTTATAAGCCGGAGCAAAGCGAGGTGACCATCATGCGTGAGATCATCAGCGTATATCGTATGCGTCTGCCGAAAGAGCGTATGCAATCGACCCGTGATAATTCCGCCCTGGGACTGGACATGTTGAATGAGCTGAAGAAGTGCCCGCGTCCGCTCTTTGATACTTTGTCTCCGGCGGACCAGAATCTGTTCCGCGTGGCAGTGACCGAGAATGAAAACACCCAGAACGACGAGACCATCGAGAACGGTATGATGCTAATGCTGCGTCGCACCGATCGTTTCCCGTCTTTGGCGTTGAGATTCATCGACACCAATCAGGTCTTCAAATCGATTCGATTTCAGGTGCATCTAGGCAACTATCGCTACAAGTTCTACGAAAAGAAATGGATTGATGGTAACGACGGTAAAGACAGAGTGCGTATTCTCCAGAAGGAGCTGAATGGTTTCGGCCGTCTGGACGAAATAGAGGTGACTCGCAACAACAAATGGGGTGACCTGATTCGCAAGATCGATCAGCCGCGTGAGGACACCTTTGAAACGGCACCGTATGTGACGGACCACCATGCAACCTACCTATTCAATAATAACCGTATCGGCCTGCTGTGGAATACCCAAGATCACATTGCGCTGTCCAACGGCATCTACCTGCCGTCCCTCTTGGAAAAGAACCGCATCCTGAGCGAAGACGAAAAACTGAAGGATCTGAAAGATCAGCACGGCAACGGTATTGCGGAATGTGTTGAACCGATGTGCTGGCTGAGCGTGTATGATATCCCAGCTGCGATCTTTCTGATCCACTTGCTTCACAAAAACGGTCAGACGTTCGAGTCTGCGGGTAAAACCGTTGAAGATATAATAAAAGCTACGTACAACAACTACCAACAATTCCTGAAGGGCATTGCGGGCGGAACCATTACCTCCATGGATGATTTACAAACCCAAGGTATCGGTATTAAGCCGTGTGATATCCCCGTCAAACTGCGTGAATATCTAAATGGCAAACAAACTAATATTAAAGAGGACTTCGTTTATCTGGGTTATGAGAGATTGCTGGACATGTATGAACGTAATGAAAAACGACTGGAGCGTTTCCAACGTGATAACGACACATACGGTAGCAAAGAAAACAAAATCGGCAAGAAGGCTTACATCGACATCCGTCCGGGTACTCTGGCTCGTCATATTGCTGATGATATCATGTTTTTTGTTCCGACGGACAAGAATAACAACAAGGTCCGGATTACCGGTCAAAACTTTAATATCATGCAAGCGGCGATTGCTCAGTACACCGGCAATATCGTGCAACTGCGTACCCTACTGCAAAAAGGCGAAATTATCAATTGTGAACGCCAGCATCCGTTCATGCACAAAGTGCTGGCGACTAAGCCGTGCAGCTTCGCAGCCCTGTACACGGCCTATCTGAAGGCGAAACGCGCATATTTGGCCAACCTCATCAAGGAATACCAAAAGGCGCGTAAAAAGACCAATGATATAAAGACCCTGAAGAAGCAGTTTGTGAAGGACTACAGCAAATTAGCGGCGCTTTCATTTCTGCACGTTGACCGCCAGAAGTGGGCGGAGCGTAACGAAGATTACTATAAACAGCTTGCCGAGAGATATGCGACGATCGAGCTGCCGCATGGTATTTTTCTGAACGCTATCCATGATGCGATTATGCAGCTGCCGGAGTCTAAGAATATGGACATTGATAAAGAAAAGGACAACGTGAGCGGCCTGATTCTGAAGTACTTCAAGGCTAGCGGTGACGCTAGTCAGCCGTTCTATGGCAACAAATTTAAGCGCAACTACAAATACTTGGACATGATGGTCAAACCGGAGTGGAGCTACTCGGGTCCGCAGCCGAGCCTGGAGAAGCACTACTATTCTATCAGCGAAATGGCGAACATGCAGAAGCAGTGGGGTCAGACCAAAGAGCTGCGCGAGAAGCGTGTTGCAGTCTATGCCAAAGCGGTGGATAAAAAAACTAACCATCTGCTGGAGAAAGCGCGTGCTAAGTTTGGGAATGACATCAGAAAGCTGGAGCGTCGTGAAAAATCGATTCAGCGTGAATGTAATGAAGAAAAAAAGGCCGCGATTGAAAAGCTCCAACATTGCCGTACCTCCTACTCCGCGAATGAGCGCGAAATCCGTCGTTACAAGGTTCAGGATATCTTGATTTTCCTAATGGCGATCGACCACCTGACCGCGGCGATGGCAAAGGTAATTTCAGCTCTTACAAGTTGCGGAACATCGGCTGCGCGAGCGAGAGCTCGGATATCCTGTCTATGCTGATGCCATTCTCTATGACGCTAGAGATTCCGAGCCTGATTAAGGACGAGCCGGCAAAGGTCATCACCATCAAGCAAGATGATTTAAAACTGAAAAACTACGGTGACTTCTTTCGTTTTATCTACGATCGTCGCATCCGTACCCTGCTGCAAAATGTTAAGGAAAACGAGTTCACCCGTCAGCAAATTGAAGAAGAACTGGAGTGCTATAATCGTATGCGGATTCCAGTGTTTAAGCAAGTTCTGTCCACCGAGGAGAAGATTTGGAAACAGTTGCCGAGTGAGCAGCTGCACAAAAAGGCGGACGACAAAGAACACCCGGTTAATATTGACTTCAAGTATTTAATGCAGTTCGTTAACATTAAAGAAAGCAATATCGAGATCATAAAAGCGATCCGCAACGCATTTAGCCATAACCAATACCCGACCGACAGCGCTGTAGCTCGTCTTGTGTTCGATAAGAAGCAGATCCCGGAGCTGGCCAAAGAGATTACCAAGTTGTTACACAACAAAACCAACCAGATTAATTTAAAAGACATGGAAGATAAGCAGTAA

***Dysgonomonas sp***

*Ds* RT-Cas1

ATGCAAACTTTATTTTGTAAACTATGCACAATCGATCACCTGTACAGCGCGTGGAATATCATCAAGACGAAGAACGCAGTTGGTGGTATTGACAAAATGACCATCCTGGATTTCGACAACAATCTTTCCGGTAATCTGAAACAGCTGCAAAAAGAGTTGCAAACCAAAGAATGGCTGCCAAATCCGTATCTGCGCATTGAGATCAAAAAAAACGAAACCGAGAAAAGGAAACTGGGCCTGTTGACCATTAAGGACAAAATCGTTCAGCAGGCGATCAAGATTTTGATTGAGCCGCGTTTTGAACGCCTGTTCCTGAATAATAGCTACGGCTATCGTCCGGATAAGGGTCATCTGAAGGCCATCAAGAGAGCTATGAGCGAAATTAACATGCGTAAAAACTTATGGGTGGCGCAGCTGGACATTGACAACTACTTCGATACCATTAACCACGAAATCTTATTTAAGCGTCTGAAGCCGATGGTTAATGATGACGAAGTACTGCGCCTGATTGACCTGTCTACGAAAATGGGTGTTGTTGATGTGAACAAGAAATGGACCGATATCACTGAGGGTGTGCCGCAGGGTGCTATACTCTCCCCGCTGTTATCTAACTTTTATCTGCACAGCTTTGATCAGTTTATTACGACCAAGACAAATTCCTATATCAGATACGCGGATGACTTCATATTTCTGAGTGAAGATGAGGAAATTATCCGTGAGCTGGTCATTAAATCGTCTACCTTCTTGAAGGAGCGTCTGTTGCTACAGTTAAACGAACCGATTATCAATGAGGTTTCCATTGGCATTGAGTTCCTGGGTATTAAGCTACAGAAAAAGAAGATCAGCATTACTGAAATCAAGCAAACCCGTTTGATCGAGTCCATCGAGATGCTGAACATTCAGCATGGCACCTTTAGCTCGAAAATTTTGCAAAAGCTTGACAGTATCAGACGTTACTATGCGCAAATCCTGCCTCAAGAGTATCTGATTCCGCTAGACAAAGCGTTGGCGGAACGCATTGTTACCCTGATCCGCACCAACACCATTGAGTTTCGTAATAAAAAGCACATCATTCAGGAGCTGAACAAGATCCCGTTTTTCAGCGCAGAAGCGAACTTGGCTCGTAAGGAGAAAATCGACGAGTGGAGCGGTCTGTATATGGACATCAAGCGTAAAAGCAATGAAAAAGACGATAACAACATCTTGGAAAAGAACAAAAAGTTGATCAATAAGCGCAAGCGCGAATACTTGAAGAAGGAGAACGAGGGCACCGAACTGGTCGTGAGCAGCTATGGCGTTTTTATTGGTAAAAGCAACAAGGGCATCATTCTAAAACAAAAGGGCAAGGTGGTGAGCGATAGCCCGAGCACCGCACTGAAGCACATCACCGTCATGACAAAAGGCGTGTCTATTAGCTCTGATGCCATCAACTACTGCATGGATAACAAGATCCCGATTGATTTCTTCGACTATACCGGTAAACACTATGCTAGCATTCTGGCACCGATCTCCGTGCAAAAAACTCTGTGGCAGATGCAGGCGAATCTGAGCAGCACGCGTAAAATCTACCTGTCCACCCAGATTATTACGGGTAAGTTAAAAAACCAGCTTAACCTGATTAAGTATTTCCACAAATACCATAAAACGGACATCGGCCTGTACCAGATTTACGAAAGCGTTAGCTCACAGCTGAGTGACATTATCAAGAAGATCAAGGACATCGCTTTGACCGAATCTAACTACAAAGAGATCCTGATGGGCTACGAGGCTCAAGGTGCCATCCTGTACTGGAACTATATTCGTCAGCTCTTACTCGACGATGGTATTGACTTCAAAAGCCGTGAGCGTAAAGGTGCCATTGACCTGTTGAACTCGCTGCTCAACTACGGCTACGCGATCATCTACGCGCGTATCTGGCAAGCGGTACTGAAAGCGAAACTGAATCCGAGCATCGGCTACCTGCACGCGTATCAACAAGGTAAACCGACCCTGGTCTATGATATCATTGAGCTGTTCCGTGCGCAAGCCGTTGATCGCATCGTGATTAGCCTGGTTCAAAAACGCGAACCACTGAGCATGTCCAACCATCTGCTTAGCGACGACACCAAAAAACTGTTGGTGCAGAATATTTTAGAACGCATTCATCGTTACGAGAAGTACCGTAGCCAAAACATTCGTTTCTCAGAAATCATCAAAGAGCAGATCCGCGATATCTCCCTTTATATCATCGGTGAAACCAATATCTTTAAACCGTATATAGCGAAATGGTAA

*Ds* Cas2

ATGAAGAGGGCTACAAAAATATTTTGTATTGTGGCGTATGATGTTGAAGACAATCGTCAGCGTGATAAGATCAGCAAACTGCTGGAAAAATATGGTATGCGTATTAACCTGTCTGTTTTCGAGTGCATGTTCACCGACACGCAATATAAAAAGGTGAAGGAGGACATCGAGAGACGCATTGATAAGCGCACTGACACCTTGGTTTACTACCCGATTTGTGTCAGCTGCTTTACCAAAATCGTGTACCAGCCGGATCGTCGCAAAAGCATTCGTACCGTAAAGATCATCTAA

*Ds* Array

TTGCGAAAATTCCTAAATGTGATGATGAGACATTCTTTCTCAGGAAGATTGGAATTGTAAATATAAACTTGATAAAATGTAGTCCTCTGAAAAACTGAATGTTGTAGAAGCTCTCACTTTGGAGGGTATGACAACATTAATCATTTCAGACGGTTTGTAGGTGTGAAAA
